# Comparative analysis of gene expression in virulent and attenuated strains of infectious bronchitis virus at sub-codon resolution

**DOI:** 10.1101/612614

**Authors:** Adam M. Dinan, Sarah Keep, Erica Bickerton, Paul Britton, Andrew E. Firth, Ian Brierley

## Abstract

Infectious bronchitis virus (IBV) is a member of the genus *Gammacoronavirus* and the causative agent of avian infectious bronchitis. IBV has a single-stranded, positive-sense RNA genome ~27 kb in length and, like all coronaviruses, produces a set of sub-genomic messenger RNAs (sgmRNAs) synthesised via the viral polymerase. Here, we used RNA sequencing (RNASeq) and ribosome profiling (RiboSeq) to delineate gene expression in the IBV M41-CK and Beau-CK strains at sub-codon resolution. Quantification of reads flanking the programmed ribosomal frameshifting (PRF) signal at the genomic RNA ORF1a/ORF1b junction revealed that PRF in IBV is highly efficient (33–40%), consistent with *in vitro* measurements. Triplet phasing of the profiling data allowed precise determination of reading frames and revealed the translation of two intergenic genes (4b and 4c on sgmRNA4), which are widely conserved across IBV isolates. RNASeq revealed two novel transcription junction sites in the attenuated Beau-CK strain, one of which would generate a sgmRNA encoding a ribosomally occupied ORF in the viral 3’ untranslated region (dORF). Within IBV transcripts, the nucleocapsid (N) protein was unexpectedly found to be inefficiently translated, despite being an abundant structural component of mature IBV virions. Finally, we demonstrate that the host cell response to IBV occurs primarily at the level of transcription, with a global up-regulation of immune-related mRNA transcripts following infection, and comparatively modest changes in the translation efficiencies of host genes.

**IMPORTANCE:** IBV is a major avian pathogen and presents a substantial economic burden to the poultry industry. Improved vaccination strategies are urgently needed to curb the global spread of this pathogen, and the development of suitable vaccine candidates will be aided by an improved understanding of IBV molecular biology. Our high-resolution data have enabled a precise study of transcription and translation in both pathogenic and attenuated forms of IBV, and expand our understanding of gammacoronaviral gene expression. We demonstrate that gene expression shows considerable intra-species variation, with single nucleotide polymorphisms associated with altered production of sgmRNA transcripts, and our RiboSeq data sets enabled us to uncover novel ribosomally occupied ORFs in both strains. We also identify numerous cellular genes and gene networks that are differentially expressed during virus infection, giving insights into the host cell reponse to IBV infection.

## INTRODUCTION

Avian infectious bronchitis virus (IBV) is a member of the genus *Gammacoronavirus* (order *Nidovirales*) and a pathogen of the domestic fowl (Cavanagh, 2005). IBV infects primarily the epithelial cells of upper and lower respiratory tract tissues, though infections can also spread to the alimentary canal, as well as to the kidneys, testes and oviduct (Cavanagh, 2007). The monopartite, polycistronic genomic RNA (gRNA) of IBV is approximately 28 kb in length, and – like those of other coronaviruses – it is 5′-methyl-capped and 3′-polyadenylated (Gorbalenya et al., 2006). Two large open reading frames (ORFs) – ORF1a and ORF1b – are situated within the 5′-proximal two-thirds of the genome. Translation of the former yields a *ca.* 3,950-aa polyprotein (pp1a); whereas translation of the latter requires −1 programmed ribosomal frameshifting (PRF) (Brierley et al., 1987; 1989) giving rise to a *ca.* 6,630-aa polyprotein (pp1ab). These polyproteins are cleaved to yield the components of the membrane-bound replication-transcription complex (RTC) (Liu et al., 1997; Brockway et al., 2003; Sawicki et al., 2007). A feature of coronavirus replication is the synthesis of a nested, 3′-coterminal set of subgenomic mRNAs (sgmRNAs) encoding the viral structural and accessory proteins. The 5′ end of each sgmRNA comprises a 56-nt sequence derived from the 5’ end of the genome, the so-called leader sequence (Brown et al., 1986; Sawicki and Sawicki, 1995). Incorporation of the leader occurs as a result of “polymerase hopping” – or discontinuous transcription – during negative-strand synthesis. When the RTC encounters specific “body transcription regulatory sequences” (TRS-Bs), the nascent negative strand can re-pair with a closely homologous leader TRS (TRS-L) at the 3′ end of the leader, after which the viral polymerase completes negative-strand synthesis using the leader as template (Fig. 1A; diamond symbols) (Brown et al., 1986; Sawicki and Sawicki, 1995, 1998; Pasternak et al., 2001; Zuniga et al., 2004; Sola et al., 2005; Sawicki et al., 2007). Subsequently, the RTC synthesises positive-strand copies of the negative-strand genomic and sgmRNAs.

**Figure 1.**
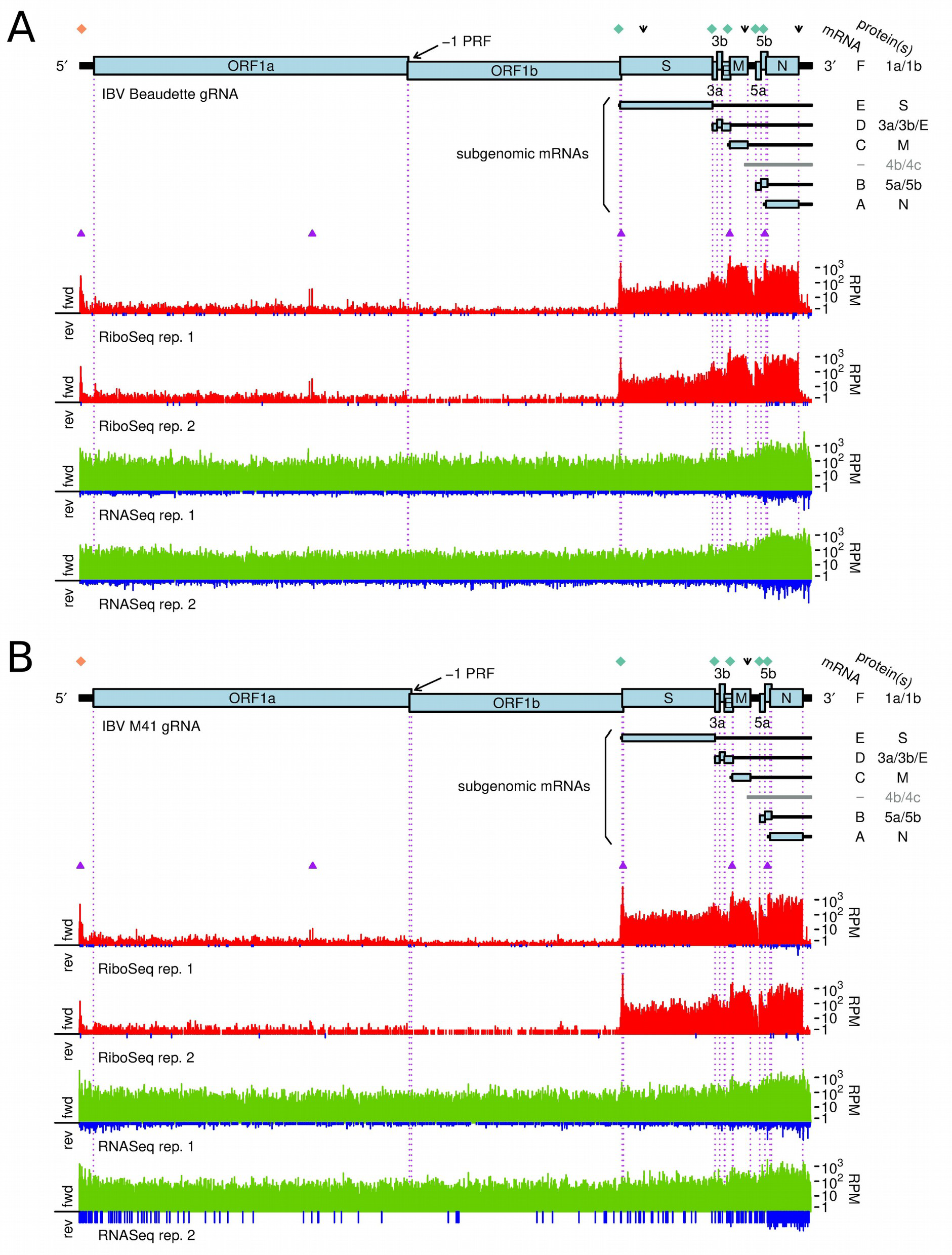
Structure and read coverage of the (A) IBV Beaudette and (B) IBV M41 genomes. Coverage in the RiboSeq (red) and RNASeq (green) libraries is plotted on a logarithmic scale. The 5′ two-thirds of the IBV gRNA contains two large ORFs encoding pp1a and pp1b, respectively. Translation of the latter requires −1 programmed ribosomal frameshifting (PRF) at the indicated site. A nested set of 3′-coterminal sgmRNAs is produced during infection. Diamond symbols show the locations of canonical TRSs at which discontinuous transcription occurs (TRS-L in orange and TRS-B in green). Downward arrows indicate the positions of non-canonical TRSs discussed in this work. Purple triangles indicate sites of ribosomal pausing (see text).

Amongst the best-characterised strains of IBV are those belonging to the Massachusetts serotype, which includes the virulent Massachusetts 41 (M41; Van Roekel et al., 1951) isolate and the laboratory-attenuated Beaudette variant (Beaudette and Hudson, 1937). Whilst M41 is restricted to growth in primary chicken cells, Beaudette is capable of replicating in both avian and non-avian cell lines; including Vero (African green monkey kidney-derived) and baby hamster kidney (BHK) cells (Cunningham et al., 1972; Otsuki et al., 1979; Alonso-Caplan et al., 1984; Casais et al., 2003). Polymorphisms in the spike (S) glycoprotein subunit 2 (S2), which spans the viral membrane, have been shown to be responsible for this variation in host cell tropism (Bickerton et al., 2018). Moreover, the S protein of M41 – but not that of Beaudette – elicits an immunoprotective response *in vivo*; although recombinant transfer of the protein from the former to the latter does not restore pathogenicity (Hodgson et al., 1984). The extent to which these strains diverge in terms of virus gene expression, or in terms of host cell gene expression in response to infection, has not been investigated in detail.

The advent of high-throughput sequencing techniques offers a means to monitor viral gene expression at unprecedented resolution (Ingolia et al., 2009; Stern-Ginossar, 2015; Irigoyen et al., 2016; 2018; Stewart et al., 2018). Here, we performed deep sequencing of ribosome-protected fragments (RPFs) – known as RiboSeq – in tandem with whole transcriptome sequencing (RNASeq), on total RNA extracts from primary chicken kidney (CK) cells infected with Beaudette and M41 strains of IBV.

## RESULTS

### RiboSeq and RNASeq Data Quality

RiboSeq and RNASeq libraries were prepared from two biological repeats each of Beau-CK-infected, M41-CK-infected, and mock-infected cells. An average of 1,156,819 RPFs and 1,727,024 RNASeq reads were mapped to viral gRNA in the virus-infected RiboSeq libraries (Supp. Table S1). The RNASeq read coverage in the library derived from the second biological repeat of M41-CK-infected cells was lower than that of other libraries due to technical losses. However, 106,741 reads were mapped to the forward strand of the viral gRNA in this case – corresponding to a coverage of approximately 3.8-fold – and these reads were generally evenly distributed along the gRNA; hence, the sequencing depth in this replicate was deemed sufficient for further analysis. The vast majority of RPFs mapping to viral and host protein-coding regions were between 27 and 29 nt in length (Supp. Fig. S1), consistent with the size of the RNA fragment protected by translating eukaryotic ribosomes from digestion by RNase I (Wolin and Walter, 1988). The length distributions of RNASeq reads were much broader, in line with the size of the gel slice excised for sequencing of fragmented RNA. RPF length was strongly related to the RPF phase relative to the reading frame of the associated coding region: 27-nt RPFs were primarily in the +1 phase; whereas 28-and 29-nt RPFs were primarily in the 0 phase (Supp. Fig. S2). As expected, RNASeq reads were far more evenly split over the three phases, with a slight bias towards phase 0 (Supp. Fig. S3), which may reflect codon usage bias – such as a preference for the use of RNY codons (Jukes, 1996; Fuchs et al., 2015; Irigoyen et al., 2016) – compounded with adaptor-ligation bias during library preparation. A meta analysis of host mRNA coding regions showed that the depth of coverage of RiboSeq 5′ read ends increased substantially 12 nt upstream of the AUG (initiation) codon for RPFs in phase 0 (generally 28- and 29-nt RPFs), and 11-nt upstream of the AUG codon for RPFs in phase +1 (generally 27-nt RPFs) (Supp. Figs. S4 and S5). This indicates that the ribosomal P-site is situated at an offset of 11 and 12 nt from the 5′ ends of RPFs for 27-nt and 28-/29-nt reads, respectively (Ingolia et al., 2011). Peaks in RNASeq 5′-read end coverage were seen at the A of initiation (AUG) codons and at the middle nucleotide of termination (UNN) codons, respectively (Supp. Figs. S6 and S7), and are considered an artefact of ligation bias.

Fig. 1 illustrates the RiboSeq (red) and RNASeq (green) read coverage of the Beaudette **(panel A)** and M41 **(panel B)** genomes. In both cases, the density of RPFs was considerably higher towards the 3′ ends of the gRNA, consistent with production of the 3′ co-terminal nested set of sgmRNAs. In contrast, RiboSeq coverage of the ORF1a and ORF1b coding sequences was relatively low; reflecting the fact that a substantial proportion of newly synthesised gRNA (but not sgmRNA) transcripts are likely to be destined for packaging rather than translation (Kuo and Masters, 2013). On average, negative-sense RNASeq reads were present at 0.28% of the level of positive-sense reads on average, indicating a ratio of positive:negative stranded RNA of ~350:1 at 24 h p.i., a ratio similar to that seen in ribosome profiling studies of the betacoronavirus mouse hepatitis virus (MHV) (Irigoyen et al., 2016). Negative-sense RPFs, which may represent contamination from ribonucleoprotein complexes (Irigoyen et al., 2016), were observed but at low abundance (0.03% of the level of positive-sense RPFs).

### Virus transcription: sequence divergence associated with IBV strain-specific TRS usage

The density of RNASeq reads mapping to a given sgmRNA represents the cumulative sum of reads derived from the gRNA and those derived from the overlapping portions of other subgenomic transcripts (Fig. 1). Therefore, to estimate the abundance of individual sgmRNAs, two independent approaches were used. First, we “decumulated” the raw RNASeq read densities mapping to inter-TRS regions, by subtracting the density of the 5′-adjacent inter-TRS region in each case (Irigoyen et al., 2016). Secondly, the abundances of chimeric RNASeq reads spanning TRS junctions were quantified, by identifying unmapped reads containing an 11-nt sequence derived from the leader region, 5′-adjacent to the TRS-L (UAGAUUUUUAA, nt 46 – 56 in Beaudette; UAGAUUUCCAA, nt 46 – 56 in M41), and including at least 16 nt 3′ of this query. Chimeric reads were assigned to specific genomic loci based on the sequences 3′ of the TRS in each case (Supp. Table S2; Fig. 2A). Overall, the chimeric read abundances for sgmRNAs were significantly correlated with the corresponding decumulated RNASeq densities (*P* < 0.01 in both cases) (Supp. Fig. S8). The sequence logos in Fig. 2B and Fig. 2C illustrate the diversity of nucleotides found at TRS-B sites identified in this study (including the novel sites discussed below) in Beaudette and M41, respectively. The core region of similarity to the TRS-L motif (CUUAACAA) is typically flanked by a 3′ adenine (A) or uracil (U) residue, and a preference for A/U residues is also seen immediately upstream of the core sequence. These flanking residues may facilitate template switching by lowering the free energy of anti-TRS-B/TRS-B duplex disassociation, since the TRS-L is also located in an AU-rich region (Sola et al., 2005).

**Figure 2.**
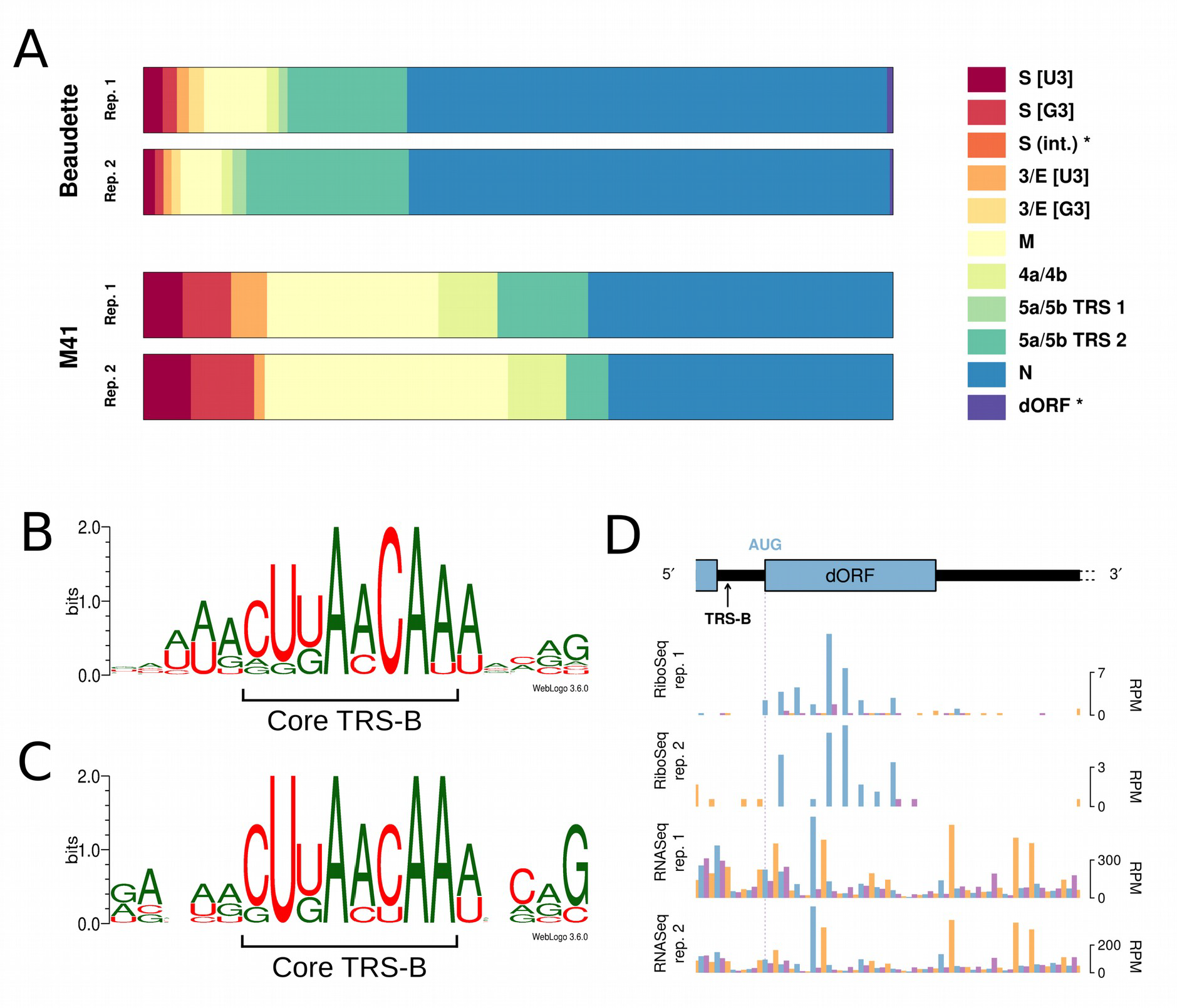
**(A)** Proportion of chimeric reads assigned to each of the indicated TRS junctions. Novel TRS identified in this study are indicated with asterisks. Note that the 5a/5b TRS 1 and the dORF TRS are present in IBV Beaudette only. **(B)** Sequence logo depicting nucleotides surrounding the identified TRS-B sites in IBV Beaudette. **(C)** Equivalent sequence logo for IBV M41. **(D)** RiboSeq and RNASeq coverage of the IBV Beaudette dORF. The location of the novel TRS-B sequence, which begins 2 nt 3′ of the N gene termination codon, is indicated with an arrow. Coverage is expressed as reads per million mapped reads (RPM). Reads in phase 0, +1, and +2 relative to the N gene ORF are shown in blue, purple, and orange, respectively; and ORFs are coloured according to the frame in which they are encoded.

Notably, the A nucleotides at positions four and seven of the core motif are the only invariant residues. In both Beaudette and M41, the TRS-B sequences associated with the S gene contain G residues at the third positions (CU<u>G</u>AACAA); in contrast to the TRS-L, which has a U at this position (CU<u>U</u>AACAA) (Supp. Table S2). Chimeric reads assigned to this gene were found to contain either a U or a G residue at position three (denoted “S [U3]” and “S [G3]”, respectively, in Fig. 2A; Supp. Table S2); with a G being more common in M41 (7.5% of reads compared with 5.8%, on average) and a U being more common in Beaudette (2.1% of reads compared with 1.5%, on average) (Supp. Table S2). These data indicate that the exact position at which discontinuous transcription occurs within a given TRS is subject to some variation, with either the TRS-L or the TRS-B templating the third residue. Similarly, the TRS-B for the 3a/3b/E genes diverges at the third position between Beaudette (CU<u>G</u>AACAA; nt 23825 – 23832) and M41 (CU<u>U</u>AACAA; nt 23832 – 23839), with the latter matching the TRS-L sequence exactly. In this case, we found that Beaudette-derived chimeric reads could contain either a U (denoted “3/E [U3]”) or a G (denoted “3/E [G3]”), with the G residue being slightly more common (1.6% versus 1.2%, respectively, on average); whereas M41-derived reads contained only the U residue (Fig. 2A; Supp. Table S2).

In agreement with a previous report (Stirrups et al., 2000), we found that the 3′-most of two adjacent canonical TRS-B sequences (both CUUAACAA; nt 25,460 – 25,467 and nt 25,471 – 25,478, labelled “5a/5b TRS 1” and “5a/5b TRS 2”, respectively, in Fig. 2A) within the 30-nt region upstream of genes 5a/5b was preferentially utilised in IBV Beaudette; accounting for 18.8% of chimeric reads on average, compared with 1.5% for the 5′-most TRS-B. Interestingly, more chimeric reads were assigned to the non-canonical TRS-B associated with genes 4b/4c (Bentley et al., 2013) – which has a low homology to the TRS-L (Supp. Table S2) – than to the first of these 5a/5b-associated TRSs in IBV Beaudette; emphasising the importance of the genomic context in facilitating discontinuous transcription (Fig. 2B and Fig. 2C) (Sola et al., 2005). Only one of the two 5a/5b TRSs (TRS 2) is found in the IBV M41 genome (Fig. 2A).

### Novel TRS in IBV Beaudette

Two additional non-canonical leader/body chimeras were identified, both specific to the Beaudette strain (Supp. Table S2). The more abundant of these (0.6% of chimeric reads) mapped to a position immediately downstream of the IBV Beaudette N gene termination codon, within the 3′ “untranslated” region (UTR). Chimeric reads derived from this site contained the sequence CUUAACAU; the last six nt of which could have been templated by the genomic (TRS-B) sequence (UAACAU, nt 27104 – 27109). There is an AUG-initiated downstream ORF (dORF) in Beaudette beginning two nt 3′ of this TRS, which comprises 11 codons (nt 27111 – nt 27143).

Inspection of our RiboSeq libraries shows that the dORF is ribosomally occupied (Fig. 2D). Such AUG-initiated dORFs are present immediately 3′ of the N genes in most IBV strains, and in TCoV, but this region appears to have been ancestrally deleted in the IBV M41 lineage; and M41 also lacks the TRS-B downstream of the N gene (UAAAAU, nt 27156 – 27161).

The second novel chimeric sequence identified in RNASeq libraries maps to a TRS-B (CUUACCAA) within the coding region of the S gene in Beaudette (nt 21242 – 21249). This is consistent with the previous detection of a sgmRNA of appropriate length via Northern blot analysis (Bentley et al., 2013). Whilst the core sequence of the TRS-B in this case is conserved in M41, there is a single nucleotide (A to C) polymorphism located four nt downstream in the 3′ flanking region, which may contribute to its lack of utilisation in this strain (Supp. Table S2).

### Virus translation: direct measurement of –1 PRF between ORF1a and ORF1b

Ribosome profiling of eukaryotic systems typically has the characteristic that mappings of the 5′ end positions of RPFs to coding sequences reflect the triplet periodicity of genetic decoding. A clear phase transition is evident in the RiboSeq libraries at the junction of ORF1a and ORF1b; where frameshifting of a proportion of ribosomes from the former ORF into the latter occurs (Fig. 3A and 3B). The mean normalised ratios of ORF1b to ORF1a RiboSeq density were 0.32 and 0.37 in IBV Beaudette and IBV M41, respectively; while the corresponding RNASeq ratios were 0.97 and 0.94, respectively (Fig. 3C). Thus on average, 33% of ribosomes in Beaudette and 39% in M41 undergo –1 PRF prior to reaching the ORF1a termination codon (Fig. 3D). These values are very similar to those measured in *in vitro* PRF assays (Brierley et al., 1987, 1989) and alongside related profiling studies of MHV (Irigoyen et al., 2016), this indicates that coronaviruses exhibit highly efficient PRF both *in vitro* and in the context of the infected cell.

**Figure 3.**
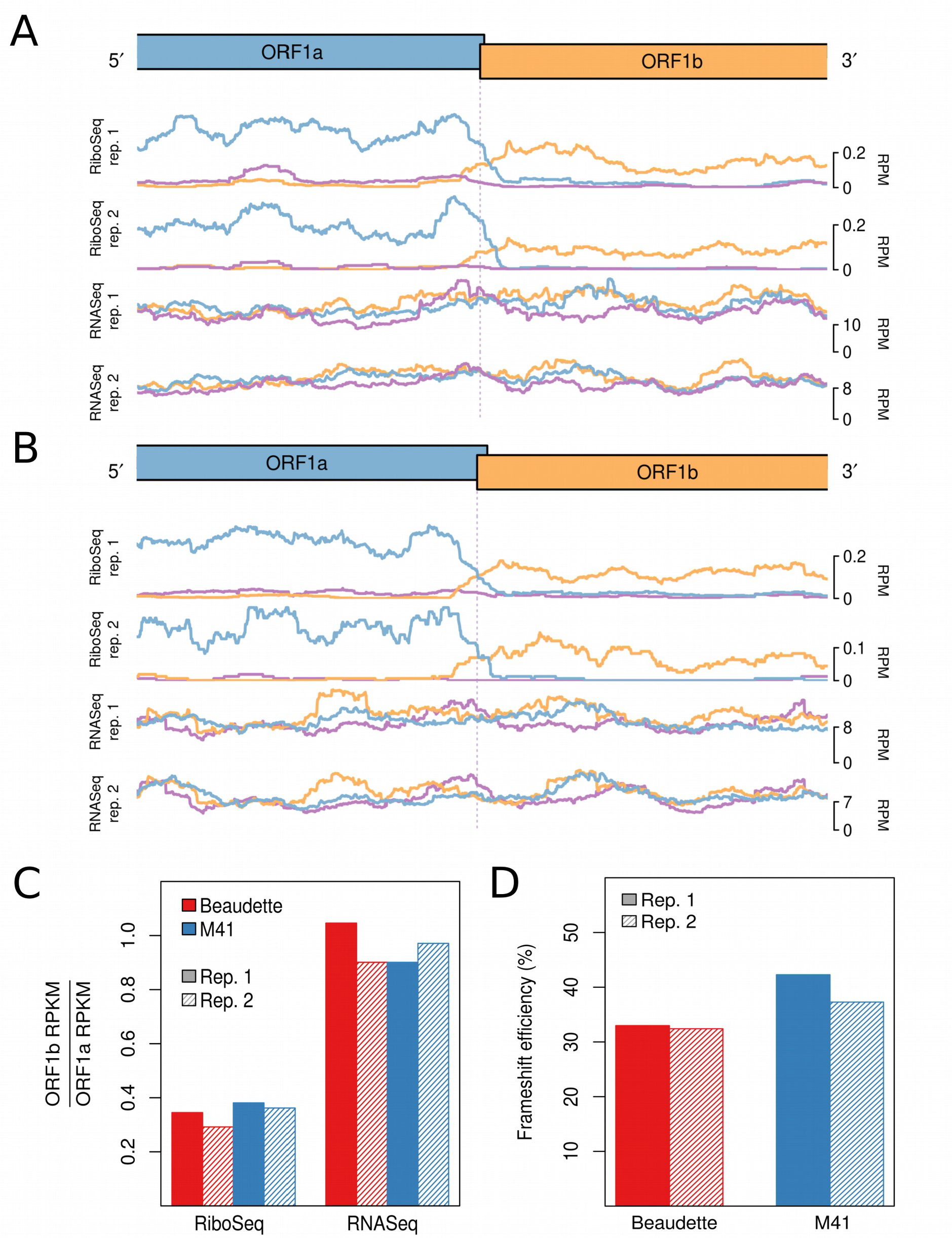
RiboSeq and RNASeq coverage proximal to the junction between ORF1a and ORF1b for IBV Beaudette **(A)** and IBV M41 **(B)**. The last 2,500 nt of ORF1a and first 2,500 nt of ORF1b are shown. Coverage is normalised to reads per million mapped reads (RPM), and smoothed with a 121-codon sliding window. Reads in phase 0, +1, and +2 relative to ORF1a are shown in blue, purple, and orange, respectively; and ORFs are coloured according to the frame in which they are encoded. **(C)** Ratios of ORF1b to ORF1a read density expressed as reads per kilobase per million mapped reads (RPKM). RPKM values exclude the 150-nt regions downstream of the ORF1a initiation codon, upstream of the ORF1b termination codon, and either side of the frameshift site. **(D)** Frameshifting efficiencies calculated using the values plotted in **(C)**.

### Ribosomal occupancy of ORF4b and ORF4c

Situated between the M and 5a genes in Beaudette and M41 is a >300-nt ostensibly “intergenic" region (IGR) (Fig. 1). No protein-coding genes are annotated here but two putative AUG-initiated ORFs are present in each virus, referred to as ORF4b and ORF4c, after their homologs in turkey coronavirus [TCoV] (Gomaa et al., 2008; Cao et al., 2008) and in the genomes of most IBV isolates (Reddy et al., 2015). The putative ORF4b genes of Beaudette and M41 are encoded by nt 25,183 – 25,335 (50 codons) and nt 25,190 – 25,474 (94 codons), respectively, of the gRNA; whereas the ORF4c genes are encoded by nt 25,339 – 25,422 (27 codons) and nt 25,395 – 25,457 (20 codons), respectively (Supp. Fig. S9). Thus, in Beaudette, the two ORFs are separated by a 3-nt spacer region and in the same reading frame (Fig. 4A); whereas in M41, ORF4c is located entirely within the ORF4b gene and in the +1 phase (Fig. 4B). Inspection of the ribosomal profiling datasets reveals substantial RPF coverage of both ORF4b and ORF4c, providing the first clear illustration that ORFs 4b and 4c are ribosomally occupied (Fig. 4). Visualisation of ORF4c translation in M41 was facilitated by good phasing in the datasets, allowing expression of both ORF4b and ORF4c to be visualised (as both blue and orange RPF peaks in the overlap region). Previous work (Bentley et al., 2013) has shown that a non-canonical TRS-B sequence – situated approximately 100 nt upstream of the M gene termination codon – facilitates production of a sgmRNA that harbours ORF4b at its 5′ end, and this TRS-B was also identified in our RNASeq data.

**Figure 4.**
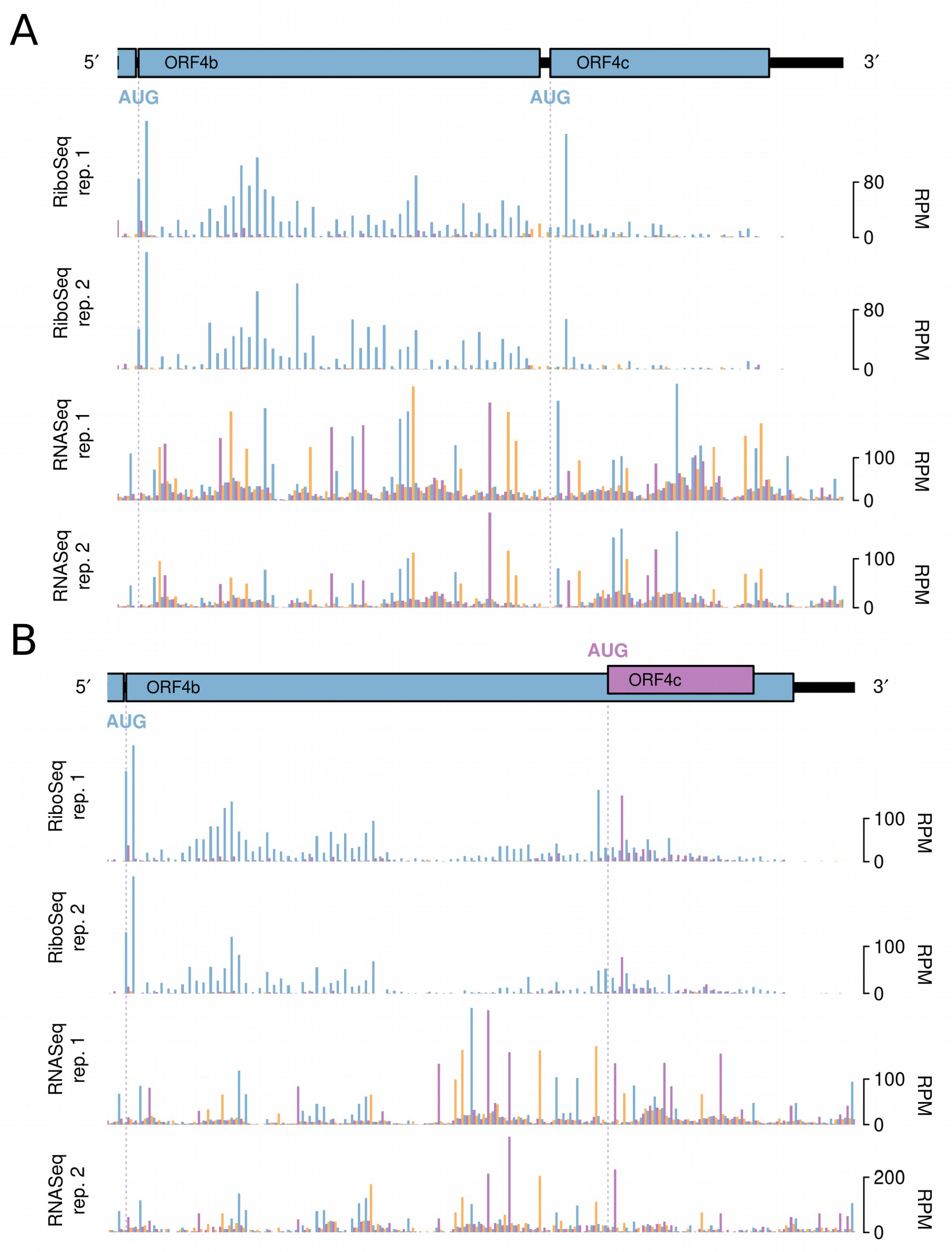
RiboSeq and RNASeq coverage of ORF4b and ORF4c in **(A)** IBV Beaudette and **(B)** IBV M41. Coverage is expressed as reads per million mapped reads (RPM). Reads in phase 0, +1, and +2 relative to ORF4b are shown in blue, purple, and orange, respectively; and ORFs are coloured according to the frame in which they are encoded.

### Translation efficiencies of IBV genes

To estimate the translational efficiency (TE) of virus genes, we summed RPFs whose 5′ end mapped in-phase between the first nucleotide of the initiation codon and 30 nt 5′ of the termination codon; thereby excluding RPFs derived from ribosomes paused during initiation or termination (Irigoyen et al., 2016). The TE of each ORF was measured as the quotient of the RPF density and the abundance of the corresponding mRNA; with separate calculations performed using the decumulated RNASeq densities and the TRS chimeric reads counts (Fig. 5 and Supp. Fig. S10, respectively). In the case of ORF4b and ORF4c, transcript abundance could not be accurately deduced via the RNASeq decumulation procedure, because the significantly lower level of expression of the 4b/4c transcript relative to that of the 5′-adjacent M gene (Fig. 1) was associated with a proportionate increase in the level of noise. Similarly, as a result of the high abundance of gRNA relative to the sgmRNA encoding S, the decumulated RNASeq density for the latter is likely to be poorly estimated, and therefore the TE value for S calculated using the chimeric read count is likely to be more accurate. From this analysis, it was observed that the 4b gene is more efficiently translated than the 4c gene; a trend also observed for the accessory genes 3a/3b and 5a/5b (Fig. 5). This is consistent with the likely requirement for leaky scanning to access the downstream ORF on each sgmRNA (see Discussion). Surprisingly, despite the fact that the nucleocapsid (N) protein is an abundant viral protein, it was not found to be efficiently translated relative to the other structural proteins, regardless of the approach used to estimate transcript abundance (Fig. 5; Supp. Fig. S10). In the case of the ORF1a and ORF1b genes, a large proportion of the genomic RNA is expected to be destined for packaging rather than translation, as mentioned above, and this probably explains the low TE values calculated for these genes (Fig. 5; Supp. Fig. S10). Additionally, the short length of the dORF precluded an accurate assessment of its translation efficiency.

**Figure 5.**
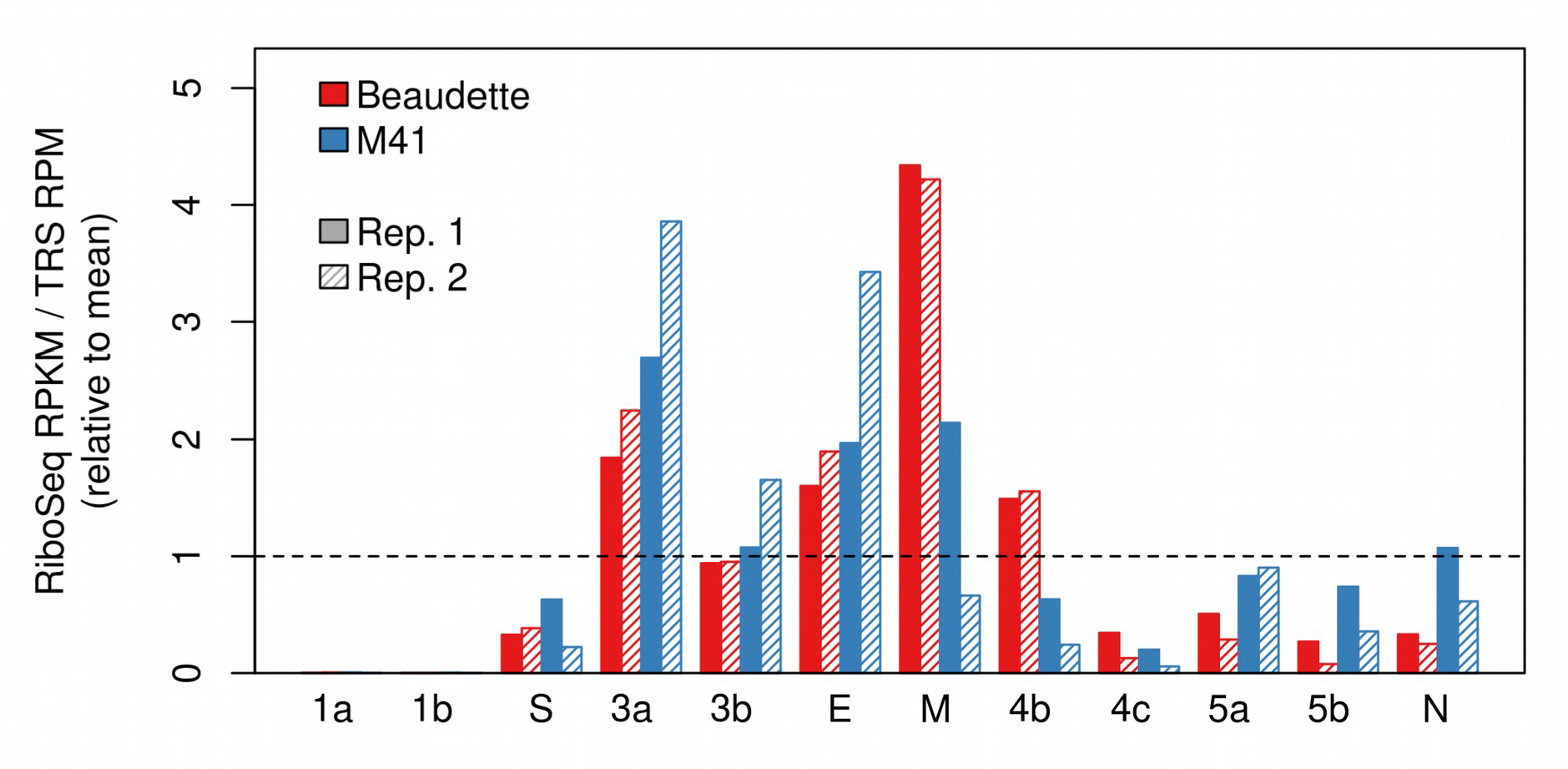
Translation efficiencies of viral genes, as calculated using the relative abundances (reads per million, RPM) of chimeric TRS-spanning RNASeq reads. Values shown are relative to the mean efficiency per TRS.

### Ribosomal pauses during IBV genome translation

Inspection of the profiling datasets revealed a number of genomic locations where RPFs accumulated to a much higher level than at neighbouring sites, indicative of ribosomal pausing. As such pauses may have biological significance, we first sought to discount those that may have arisen artefactually. The known translation initiation sites in the virus genome generally showed high ribosome occupancy, but as the infected cells were treated with cycloheximide (CHX) prior to lysis to “freeze” ribosomes onto the mRNA, these pauses are likely to be over-represented, as ribosomes can accumulate at start codons during the CHX treatment period (Ingolia et al., 2011). Fluctuations in RPF density can also occur as a result of nuclease, ligation, and PCR biases during library construction. As the latter two biases can also occur during RNASeq library generation, we also discounted any pauses that had an obvious counterpart in RNASeq datasets. With these criteria, we identified five obvious sites of ribosomal pausing conserved in Beaudette and M41, one in the 5′ UTR and four within the coding region (indicated in Fig. 1, purple triangles; see Table 1). Pauses in 5′ UTRs can represent ribosomes initiating at upstream ORFs (uORFs), although in both Beaudette and M41, the P-site of the ribosome paused over bases 28–56 in the 5′ UTR of the genome is on a non-AUG codon (UUG) in a weak Kozak initiation consensus. As this pause is located upstream of the TRS_L, it reflects the sum of pausing on all sgmRNAs. To view the extent of the pause in context, we remapped reads to the most abundant sgmRNA, i.e. that of the N gene (Fig. 6). As can be seen, the “Leader pause” remains clearly evident (as is a smaller pause three codons downstream), albeit smaller in magnitude than those pauses seen at an N uORF (see below) and the authentic AUG codon of the N protein. Initiation at the UUG codon would result in translation of solely a dipeptide and thus the pause, if biologically relevant, may act as a regulator of downstream initiation events rather than through the encoded product. We note that an equivalent Leader pause is seen in MHV (UUG codon, 1-codon ORF; Irigoyen et al., 2016). It is possible that pausing at this codon is potentiated by queueing of initiating ribosomes on sgmRNAs. The origin of the pauses within the coding region are enigmatic. The two adjacent pauses referred to collectively as Pause 2 in Table 1 correspond to translation of a region of non-structural protein 4 (nsP4) downstream of the membrane spanning domains (Oostra et al., 2007; Doyle et al., 2018). It is feasible that ribosomes pause here whilst the nascent peptide is being folded into membranes. Pauses 3 and 4 are noticeably large and correspond to ribosomes pausing soon after initiation of the S and M proteins, respectively. In the case of the former, the pause appears to be unrelated to the signal sequence at the N-terminus of the S protein, since this would still be within the peptide exit tunnel of paused ribosomes. Pause 5 corresponds to a potential non-AUG uORF (AUU, in a reasonable context) within the N mRNA (Fig. 6).

**Table 1.**
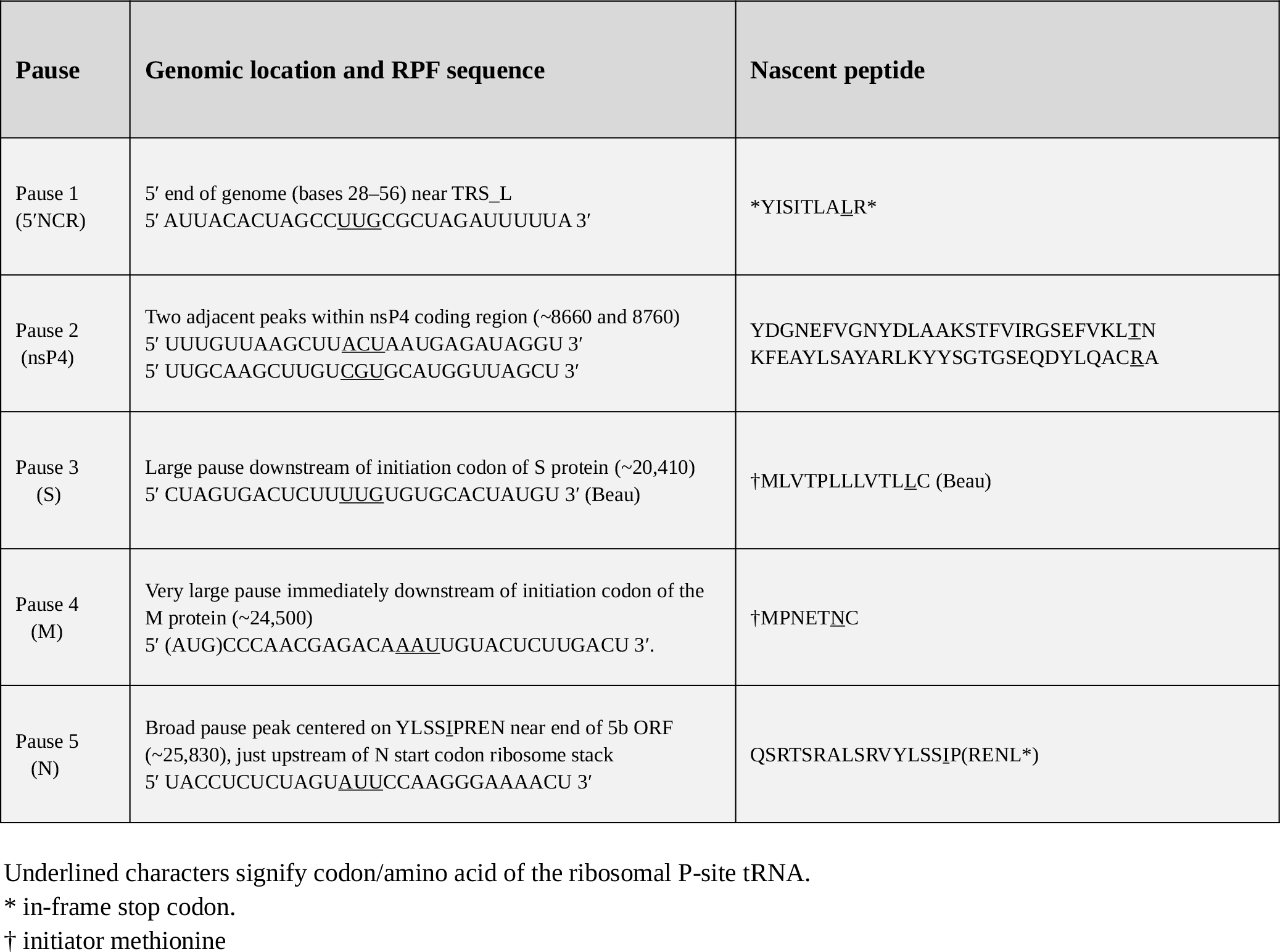
Ribosomal pause sites within the IBV genome.

**Figure 6.**
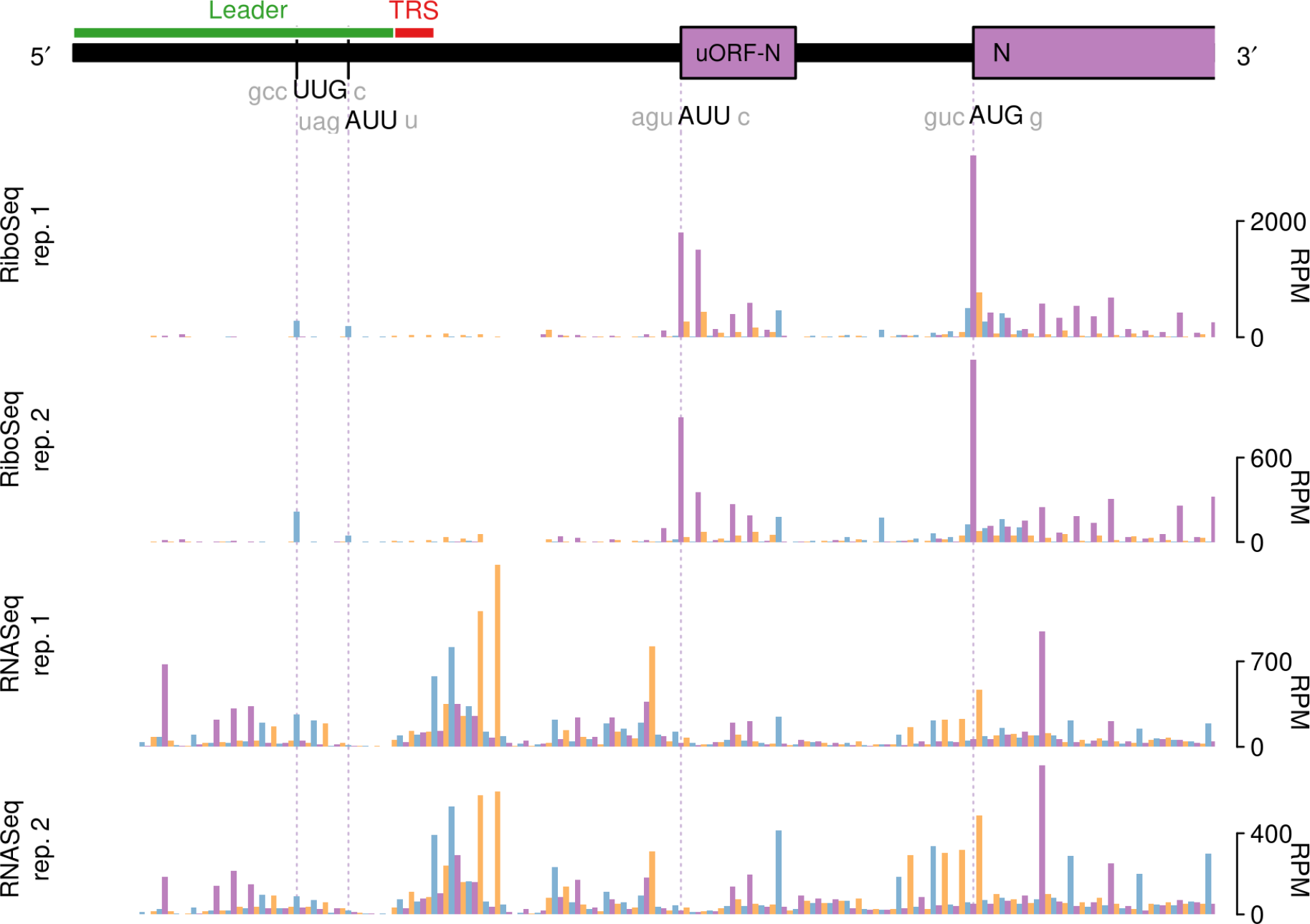
RiboSeq and RNASeq coverage of sgmRNA N in IBV Beaudette. Coverage is expressed as reads per million mapped reads (RPM). Reads in phase 0, +1, and +2 relative to N are shown in purple, orange and blue, respectively; and ORFs are coloured according to the frame in which they are encoded.

It is noteworthy that in our analysis of ribosomal pause sites, we did not see pausing at the AUG of the previously described 11 amino acid uORF of the genomic mRNA (AUG at nt 131–133; Boursnell et al., 1987), and indeed there were few reads on the uORF itself, indicating that it is not heavily translated. Further, no pausing was observed at the PRF site at the ORF1a/ORF1b overlap.

### Differential expression of host genes in response to IBV infection

We investigated the differential transcription and translation of host genes in response to IBV infection by comparing RNA and RPF densities per coding region for infected samples and mocks (see Materials and Methods). Details of the genes found to be differentially expressed (DEGs) (FDR < 0.05 with multiple testing correction using the Benjamini-Hochberg method) at the level of transcription (4,266 genes) or translation (3,627 genes) respectively, are provided in **Supp. Data Sets S1 and S2.** Overall, the patterns of change in host cell gene expression in response to infection were broadly similar for Beaudette and M41, with positive inter-strain correlations in the log2 fold changes (log2FC) in transcript abundance and translation efficiency (R^2^ = 0.95 and R^2^ = 0.85, respectively, *P* values both < 2.2 × 10^−16^; Fig. 7A). Notably, the majority of differentially transcribed genes were up-regulated rather than down-regulated (i.e. log2FC > 0) for both strains (Fig. 7A; **left panel**), with 2.1-fold and 3.5-fold more up-regulated compared with down-regulated transcripts (FDR <0.05; see Materials and Methods) detected in Beau-CK-infected cells and M41-CK-infected cells, respectively (**Supp. Data Set S1**). This effect was not seen at the level of translation, where there were fewer differentially expressed genes overall, and the logFC values of those genes were more evenly distributed around 0, with slight skewing towards negative values (i.e. reduced TE) (Fig. 7A; **right panel** and **Supp. Data Set S2**). The core host transcriptional response to the two strains involved 579 commonly up-regulated and 132 commonly down-regulated genes, while the core translational response consisted of 34 commonly up-regulated and 79 commonly down-regulated genes. Gene ontology (GO) term enrichment revealed that numerous immune-related pathways were among the most significantly enriched terms in the core response sets (Fig. 7B and Fig. 7C). There was also evidence of integration and coordination of responses at the transcriptional and translational levels. For example, the GO term “positive regulation of NF-kappaB transcription factor activity” (GO:0051092) was enriched among transcriptionally up-regulated genes; whereas “negative regulation of NF-kappaB transcription factor activity” (GO:0032088) was enriched among translationally down-regulated genes. In a direct inter-strain comparison of statistically significant DEGs we identified 51 differentially transcribed genes, 45 of which were more highly expressed in Beaudette-infected samples, and six of which were more highly expressed in M41-infected samples (**Supp. Data Set S1**). The most significantly enriched GO term in the former set was “regulation of signaling receptor activity” (GO:0010469); while pro-proliferative and anti-apoptotic GO terms were also enriched (Supp. Table S3). The latter set included three heat shock protein-encoding genes, and consequently the top enriched GO terms were related to “protein refolding” (Supp. Table S4). Just one gene (ENSGALG00000015358 [MYH15], encoding myosin heavy chain 15) had a significantly higher translation efficiency in M41-infected samples compared with Beaudette-infected samples.

**Figure 7.**
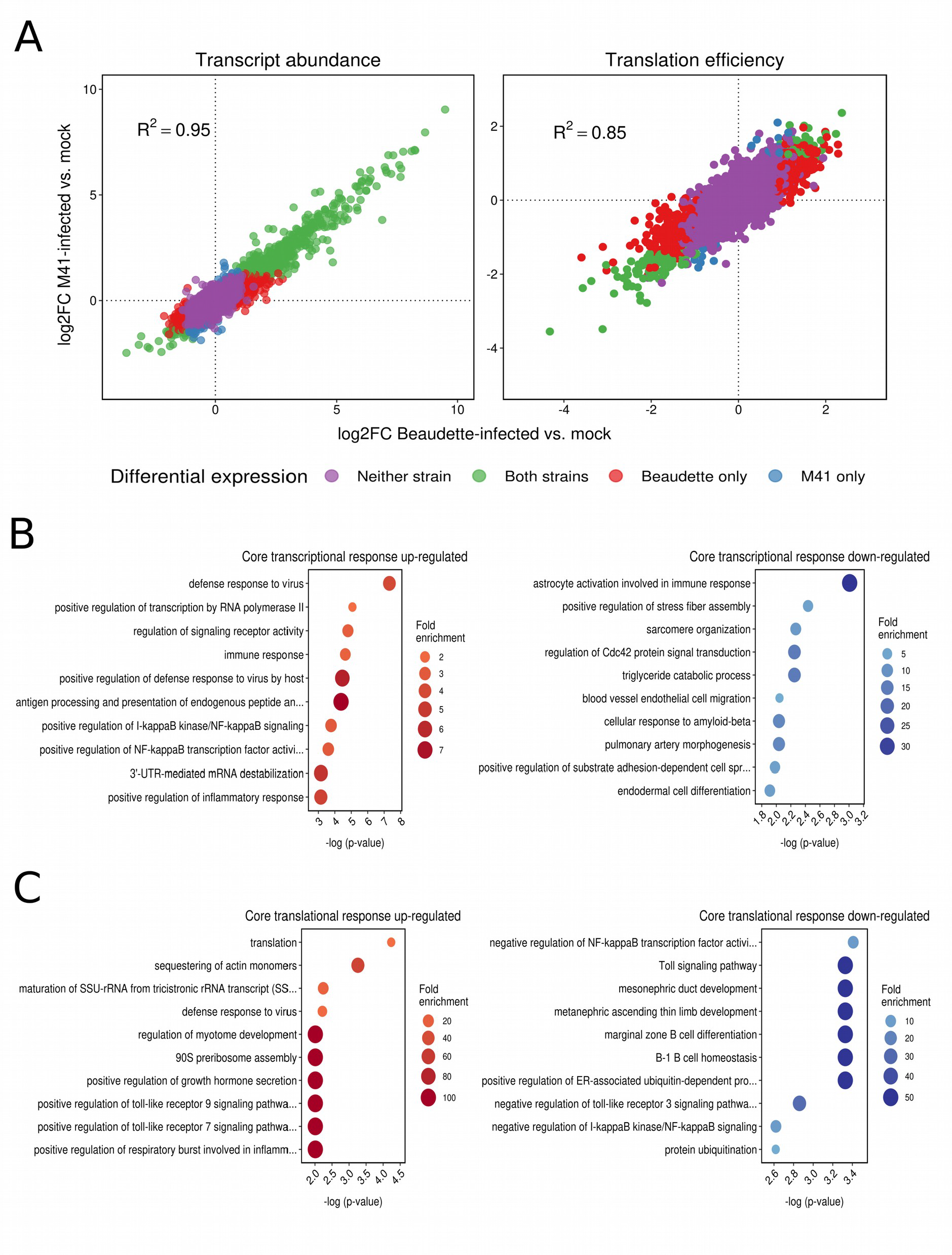
**(A)** Log2 fold changes (log2FC) in host transcript abundance and translation efficiency in infected cells relative to mocks. In both cases, a high degree of correlation was observed between logFC values in Beaudette-infected samples (*x*-axes) and M41-infected samples (*y*-axes), with transcript abundances skewed towards positive log2FC values. **(B)** The ten most significantly enriched GO terms among commonly up-regulated (left panel) and commonly down-regulated (right panel) genes at the level of transcription. **(C)** The ten most significantly enriched GO terms among commonly up-regulated (left panel) and commonly down-regulated (right panel) genes at the level of translation efficiency.

In comparisons of host gene expression between Beaudette-, M41- and mock-infected cells, the significantly differentially expressed genes (FDR <0.05) were ranked by log2FC (**Supp. Data Set S3**) and the top 100 DEGs (or fewer) within each category were subjected to STRING analysis (Szklarczyk et al., 2017) to identify potential protein-protein interaction pathways (Fig. 8 and Supp. Fig. S11). A selection of the key pathways proposed and examples of the associated genes are shown in Table 2. Clear patterns of host response to virus infection were present that are discussed below. Note in inter-strain comparisons of M41 versus Beaudette, only the transcriptionally downregulated category had sufficient gene candidates for STRING analysis; the other three categories had a total of only seven DEGs (thus no plots are shown in the **Supp. Info.** for these).

**Table 2.**
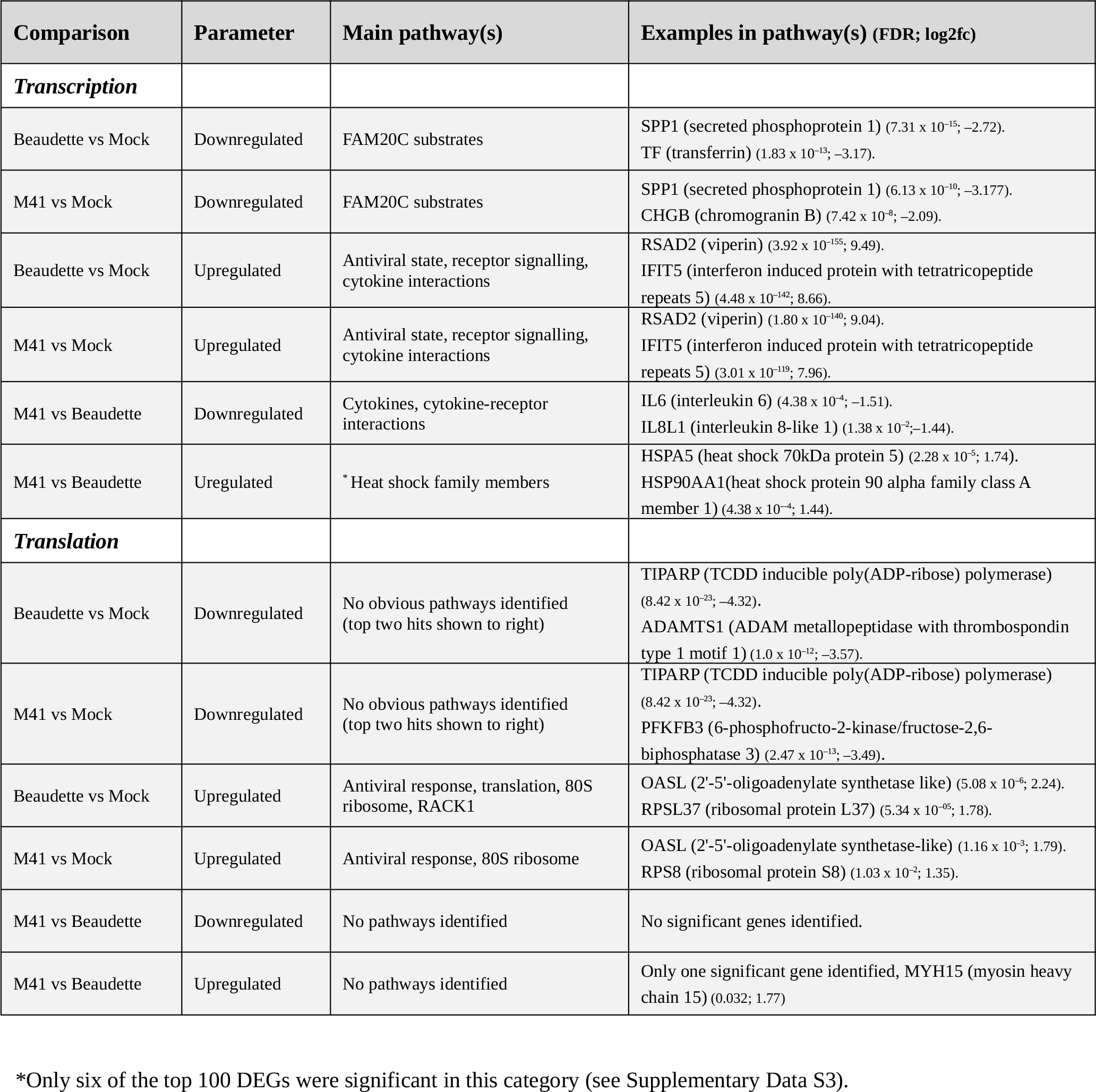
STRING analysis of differential gene expression.

**Figure 8.**
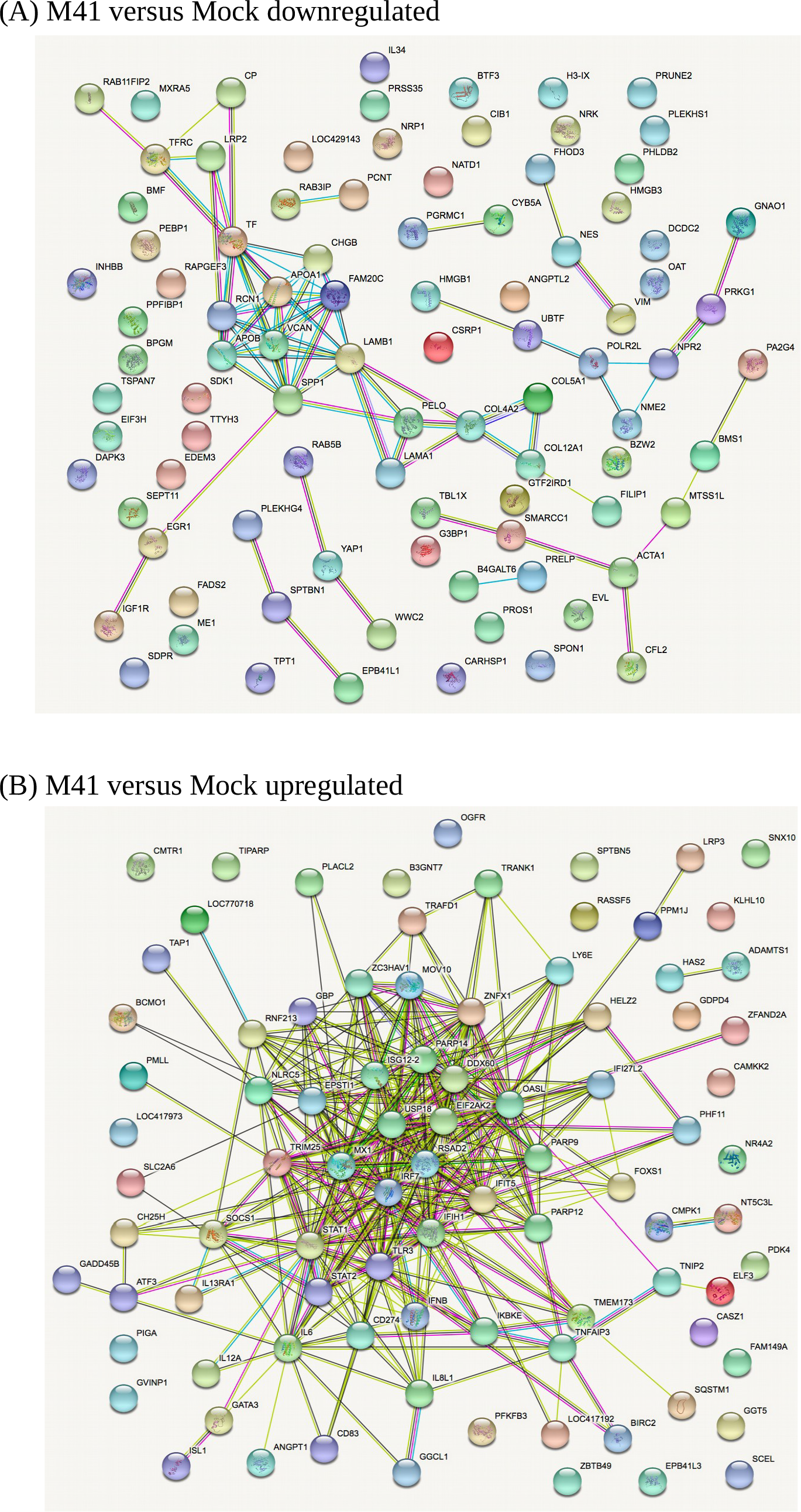
STRING analysis of the relationship between differentially expressed transcripts in comparisons of IBV M41- and mock-infected cells. **(A)** Downregulated genes **(B)** Upregulated genes. The network nodes represent the proteins encoded by the differentially expressed genes. Seven different coloured lines link a number of nodes and represent seven types of evidence used in predicting associations. A red line indicates the presence of fusion evidence; a green line represents neighborhood evidence; a blue line represents co-ocurrence evidence; a purple line represents experimental evidence; a yellow line represents text-mining evidence; a light blue line represents database evidence; and a black line represents co-expression evidence.

## Discussion

Here, we describe the first high-resolution study of gammacoronaviral gene expression during infection of primary chick kidney cells. Analysis of RNASeq data sets through chimeric read analysis or decumulation allowed us to quantify the relative levels of viral genomic and subgenomic mRNAs and to define the sequence diversity of strain-specific TRS utilisation. The predominant sgmRNA in both strains was that encoding the N protein, and between strains, the M transcript was relatively more abundant in M41. In Beaudette, two novel TRS were identified, one in the viral 3′ UTR immediately downstream of the N gene termination codon, and one mapping to a TRS-B within the S gene. In the former, a short ORF (dORF) – initiated two nt 3′ of the TRS – is present and ribosomally occupied. The potential biological relevance of this ORF remains to be determined; such dORFs are present in most IBV strains and in TCoV, but it is lacking in M41 (as is the TRS-B). A recent report has described the same sgmRNA (initiating at the identical TRS) as a novel non-coding RNA of IBV (An et al., 2019). The S gene TRS-B, proposed earlier (Bentley et al., 2013), was also identified.

RiboSeq analysis, in conjunction with RNASeq, revealed that the N protein is not more efficiently translated than other structural proteins, despite being a structural component of IBV virions. Coronaviral N proteins are highly basic, reflecting the fact that they associate tightly with the negatively charged gRNA (Laude et al., 1995), and a previous meta-analysis has indicated that the decoding of positively charged amino acid residues is associated with a lowering in the speed of translation (Charneski and Hurst 2013; Sabi & Tuller, 2015; Requiao et al., 2016). Thus, it is possible that the relatively low TE of N is an unavoidable consequence of its amino acid composition. N expression may also be regulated by a putative uORF whose initiation codon is located some 50 nt upstream of the N AUG codon (Fig. 6).

The efficiency of PRF at the IBV ORF1a/ORF1b overlap in natural infection was found to be 33– 40%. This range is in close agreement with previous *in vitro* measurements of IBV frameshifting efficiency carried out using reporter constructs (Brierley et al., 1989) and is consistent with the notion that frameshifting in IBV is among the more efficient examples of canonical eukaryotic −1 PRF signals that have been studied to date (Atkins et al., 2016; Irigoyen et al., 2016). Whether the modest difference in –1 PRF efficiency measured for Beau-CK and M41-CK has biological significance is uncertain, and it may represent experimental variation. The frameshift signal of M41-CK differs from Beau-CK in only three of 81 nucleotide positions, all of which are located in loop 3 of the stimulatory pseudoknot and not expected to affect pseudoknot function or stability (Brierley and Pennell, 2001). As also described for MHV-infected cells (Irigoyen et al., 2016), there was no evidence that the frameshift-stimulatory pseudoknot induced ribosomal pausing on the slippery sequence. Thus pausing may not be a component of the frameshifting mechanism, or the pause may be too short-lived to be visualised by the profiling technique.

A meta-analysis of host genes revealed highly specific phasing of the RiboSeq data sets, enabling the accurate determination of the reading frame of translation for individual RPFs. Good phasing in the datasets and substantial read depth also allowed us to examine translation of viral accessory ORFs. It was evident that both 4b and 4c are efficiently translated, at levels comparable to those of the 5a/5b accessory protein-encoding genes. The mechanism by which ribosomes might access ORF4c, however, is not clear. Given the absence of AUG codons within the regions between the 5′ ends of ORF4b and ORF4c in both Beaudette and M41 (Supp. Fig. S9), and the weak initiation context of the ORF4b start codon, it is possible that a proportion of ribosomes might bypass the ORF4b initiation codon and instead translate ORF4c via “leaky scanning” (Firth and Brierley, 2012), although it should be noted that intervening AUG codons do exist in some other IBV strains (Supp. Fig. S9).

The relevance to virus gene expression of the sites of significant ribosomal pausing identified in the genome remains to be investigated experimentally. Two of these pause sites appear to correspond to uORFs initiated at non-AUG initiation codons, one in the 5′ UTR and one upstream of the N gene. In each case, ribosomes initiating on the main ORF AUG (ORF1a and N respectively) could potentiate initiation on the uORFs through stacking of scanning ribosomes, and this could be artefactually increased by the cycloheximide pretreatment used during sample preparation. Two of the other pause sites correspond to ribosomes paused post-initiation early in the coding regions of the S and M genes. A biological explanation for this is lacking at present. We are aware that the treatment of yeast cells with cycloheximide can lead to an early block in elongation in stressed cells (Gerashchenko and Gladyshev, 2014; Duncan and Mata, 2017), but meta-analysis of host genes in our infected cells does not reveal an obvious elongation block. Further, the S and M genes show deep ribosome coverage along their lengths, inconsistent with a block in elongation. The remaining pause site (in fact a doublet) appears during translation of the nsP4 region of the polyprotein close to the C-terminus of nsP4. Coronaviral nsP4 proteins are important for the membrane rearrangements required for viral RNA synthesis and contain multiple membrane spanning domains (Doyle et al., 2018). A possible explanation for the ribosomal pauses seen here is that translation is paused to permit the correct folding of nsP4 into membranes.

In general, the pauses we discern during translation of the IBV genome are discrete, substantial in terms of read counts, and reproducible. As mentioned above, their origin is uncertain, but it seems unrelated to the identity of the P-site tRNA. Recent studies have shown that P-site prolyl-tRNA is a strong determinant of ribosomal pausing, partly due to the slow rate of peptide bond formation with this amino acid (Sabi and Tuller, 2015; 2017). However, none of the stall sites identified here have proline tRNA in the P-site. One possible explanation we cannot rule out is that some feature of the nascent peptide engenders pausing through interactions with the ribosome exit tunnel or chaperones; clusters of positively-charged amino acid residues have been documented to induce ribosome pausing (Ingolia, 2014; Sabi and Tuller, 2017), although such clusters are not evident upstream of the pause sites seen here. The recent developments of methodologies and algorithms to identify and characterise ribosomal pause sites may clarify the situation in future (Chadani et al., 2016; Kumari et al., 2018).

Our data indicate that the host response to IBV is mediated primarily at the level of transcription, with the up-regulation of hundreds of genes, many of which have immune-related functions. Changes in translational efficiency were more modest, with more genes showing decreased rather than increased translation in response to IBV infection. Many of the transcriptionally upregulated genes identified reflect the host response to virus infection, as seen previously with IBV infection of chickens (Cong et al., 2013; Chhabra et al., 2018) and with other RNA viruses (Zhang et al., 2018). Some of the protein pathways identified have not been associated with coronavirus infection previously and warrant experimental follow up, for example, the potential downregulation of transcription of genes linked to FAM20C, a kinase that generates the majority of the extracellular phosphoproteome (Tagliabracchi et al., 2015). Also of interest is the translational upregulation of ribosomal protein synthesis in infected cells for both Beau-CK and M41-CK. A direct comparison of DEGs in M41-CK and Beau-CK-infected primary chick kidney cells did not identify any obvious pathways that would reflect their differential pathogenesis. Overall, these data contribute towards a substantially improved understanding of the early innate immune response to IBV infection, including distinct features of the transcriptional and translational responses.

## MATERIALS AND METHODS

### Virus and cells

The apathogenic molecular clone of IBV, Beau-R, has been described previously (Casais et al., 2001) and was used to generate virus Beau-CK. The pathogenic isolate M41-CK has been described previously (Kottier et al., 1995). Primary chick kidney (CK) cells were produced from 2–3 week-old specific pathogen free (SPF) Rhode Island Red chickens (Hennion and Hill, 2015). CK cells (0.8 × 10^6^ cells/ml) were plated in 10-cm dishes and upon reaching 100% confluence (two days post-seeding) were washed once with PBS and infected with 9.6 × 10^6^ PFU Beau-CK or M41-CK. After 1 hour incubation at 37 °C, 5% CO_2_, the inoculum was removed and replaced with 10 ml fresh 1× BES (1X minimal essential Eagle’s medium [MEM], 0.3% tryptose phosphate broth, 0.2% bovine serum albumin, 20 mM N,N-Bis(2-hydroxyethyl)-2-aminoethanesulfonic acid (BES), 0.21% sodium bicarbonate, 2 mM L-glutamine, 250 U/ml nystatin, 100 U/ml penicillin, and 100 U/ml streptomycin). Cells were harvested at 24 hours post-infection when clear regions of cytopathic effect (CPE) were visible.

### Drug treatment, cell harvesting and lysis

Cycloheximide (CHX; Sigma-Aldrich) was added directly to the growth medium (to 100 µg/ml) and the cells incubated for 2 min at 37 °C before rinsing with 5 ml of ice-cold PBS containing CHX (100 µg/ml). Subsequently, dishes were incubated on ice and 400 µl of lysis buffer [20 mM Tris-HCl pH 7.5, 150 mM NaCl, 5 mM MgCl_2_, 1 mM DTT, 1% Triton X-100, 100 µg/ml cycloheximide and 25 U/ml TURBO™ DNase (Life Technologies)] dripped onto the cells. The cells were scraped extensively to ensure lysis, collected and triturated with a 26-G needle ten times. Lysates were clarified by centrifugation for 20 min at 13,000 g at 4 °C, the supernatants recovered and stored at −80 °C.

### Ribosomal profiling and RNASeq

Cell lysates were subjected to RiboSeq and RNASeq. The methodologies employed were based on the original protocols of Ingolia and colleagues (Ingolia et al., 2009; 2012), except ribosomal RNA contamination was removed using a commercial RiboZero Gold magnetic kit (Illumina) and library amplicons were constructed using a small RNA cloning strategy (Guo et al., 2010) adapted to Illumina smallRNA v2 to allow multiplexing. The methods used were as described by Chung et al. (2015), except the 5′ and 3′ adapters included seven consecutive randomised bases at the 3′ and 5′ ends (respectively). This facilitated removal of reads duplicated during polymerase chain reaction (PCR) amplification of cDNA libraries (Aird et al; 2011) and reduced ligation bias. Amplicon libraries were deep sequenced using an Illumina NextSeq platform (Department of Pathology, University of Cambridge).

### Computational analysis of RiboSeq and RNASeq data

Adaptor sequences were trimmed using FASTX-Toolkit (hannonlab.cshl.edu/fastx_toolkit), and reads shorter than 25 nt following adaptor trimming were discarded. Mapping was performed using Bowtie version 1 (Langmead et al., 2009) with parameters -v 2--best (i.e. maximum 2 mismatches, report best match). Adaptor-trimmed, de-duplicated reads were mapped sequentially to host (*Gallus gallus*) ribosomal RNA (rRNA); IBV Beaudette (GenBank accession: NC_001451.1) or IBV M41 (GenBank accession: DQ834384.1) gRNA; Ensembl host non-coding RNA (ncRNA); NCBI RefSeq host mRNA; and to the host genome. The order of mapping was tested to check that virus-derived reads were not lost accidentally due to mis-mapping to host RNA, or *vice versa*. When performing analyses of viral and host gene expression, only 28- and 29-nt RiboSeq reads (corresponding to RPFs mapping primarily in phase 0) and only ≥ 40 nt RNASeq reads were used. A 12-nt offset was applied to the 5′ mapping positions of RPFs, to approximate the P-site position of the ribosome (see Supp. Fig. S4 and Irigoyen et al., 2016). To normalize for different library sizes, reads per million mapped reads (RPM) values were calculated using the sum of total virus RNA plus total host RefSeq mRNA (positive sense reads only) as the denominator.

Host mRNA RiboSeq and RNASeq phasing distributions were derived from reads mapping internally to the coding regions of ORFs; specifically, the 5′ end of the read had to map between the first nucleotide of the initiation codon and 30 nt 5′ of the last nucleotide of the termination codon, thus, in general, excluding RPFs of initiating or terminating ribosomes. Histograms of 5′ end positions of host mRNA reads relative to initiation and termination codons (Supp. Figs. 4 – 7) were derived from reads mapping to RefSeq mRNAs with annotated CDSs at least 450 nt in length and annotated 5′ and 3′ UTRs at least 60 nt in length. When calculating the translation efficiencies of viral genes, only in-phase (i.e. phase 0 with respect to the ORF in question) RiboSeq reads were counted.

For host differential expression analyses, non-ribosomal, non-viral reads in each library were mapped to the *Gallus gallus* 5.0 assembly (December 2015) using STAR (Dobin et al., 2013), with gene annotations from Ensembl release 94 (Cunningham et al., 2019). A maximum of two mismatches were allowed when mapping. Read counts per gene (protein-coding genes only) were obtained using HTSeq (Anders et al., 2015), with a requirement that reads map entirely within the forward strand coding sequence (htseq-count parameters: -m intersection-strict -s yes -t CDS). For each comparison of experimental groups, only genes with an average of at least 50 mapped reads were included in differential expression analyses. GO term enrichment analysis was carried out using the topGO package in R (Alexa and Rahnenfuhrer, 2018) and Fisher’s exact test was used to assess the enrichment of individual GO terms in specific gene lists. Protein-protein interaction networks were constructed using the Search Tool for Retrieval of Interacting Genes (STRING) database (Szklarczyk et al., 2017).

## Acknowledgements

This work was supported by UK Medical Research Council [MR/M011747/1] and Wellcome Trust [202797/Z/16/Z] grants to I.B. and Wellcome Trust grant [106207] and European Research Council (ERC) European Union’s Horizon 2020 research and innovation programme grant [646891] to A.E.F.

## DATA AVAILABILITY

Sequencing data have been deposited in ArrayExpress (http://www.ebi.ac.uk/arrayexpress) under the accession number E-MTAB-7849.

---

**Table S1.** Library composition statistics: table of read counts per library. Random 7nt sequences were added to the ends of both 5′ and 3′ adaptors during library preparation to facilitate the removal of duplicate reads introduced during PCR. All libraries were de-duplicated following the removal of the adaptor sequences, as described in the Materials and Methods section. Numbers of reads mapped to the forward (fwd) and reverse (rev) strands of the viral gRNA are shown separately in each case; only forward strand-mapping reads are included in the host mRNA counts.

| Type | Condition | Total | De-duplicated | viral gRNA |  | host mRNA |
| --- | --- | --- | --- | --- | --- | --- |
|  |  |  |  | fwd | rev |  |
| Repeat 1 |  |  |  |  |  |  |
| RiboSeq | Beaudette-infected | 55,648,104 | 32,461,160 | 1,470,253 | 630 | 3,070,823 |
|  | M41-infected | 47,459,412 | 24,416,659 | 1,838,613 | 792 | 4,193,904 |
|  | Mock-infected | 46,640,167 | 26,170,072 | 399 | 1 | 9,400,391 |
| RNASeq | Beaudette-infected | 27,720,370 | 12,171,506 | 2,365,784 | 4,842 | 882,421 |
|  | M41-infected | 24,939,168 | 14,070,349 | 1,829,331 | 4,070 | 1,616,990 |
|  | Mock-infected | 25,036,626 | 15,462,396 | 2,737 | 9 | 2,037,543 |
| Repeat 2 |  |  |  |  |  |  |
| RiboSeq | Beaudette-infected | 55,392,595 | 30,089,677 | 778,394 | 191 | 3,017,330 |
|  | M41-infected | 51,319,886 | 27,664,513 | 540,016 | 63 | 1,978,766 |
|  | Mock-infected | 56,813,327 | 26,535,892 | 159 | 1 | 3,413,589 |
| RNASeq | Beaudette-infected | 26,792,717 | 15,637,095 | 2,606,240 | 8,245 | 1,840,748 |
|  | M41-infected | 46,163,533 | 2,680,836 | 106,741 | 388 | 127,579 |
|  | Mock-infected | 26,095,361 | 15,802,166 | 4,589 | 8 | 1,920,949 |

**Table S2.**
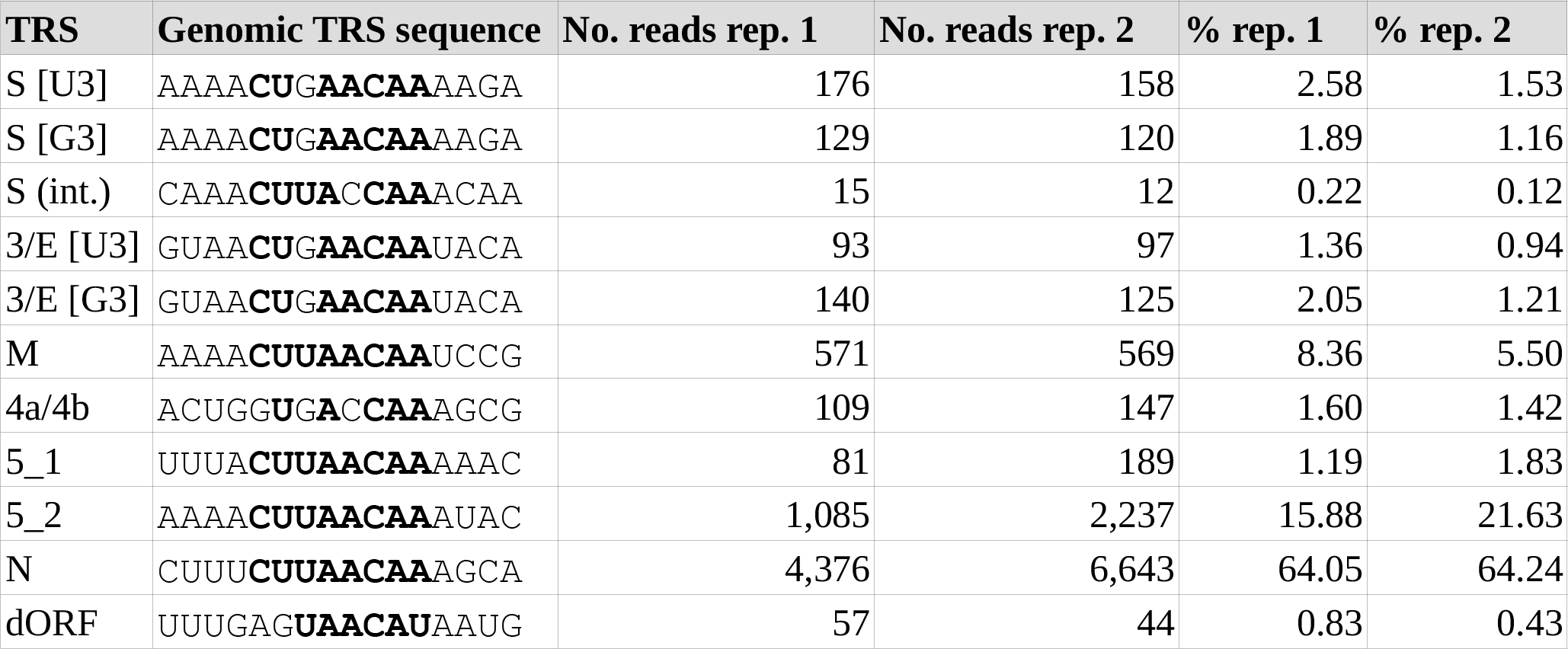
**(A)** Chimeric TRS-spanning reads identified in IBV Beaudette samples. Residues matching the leader TRS (TRS-L) are shown in bold.

**Table S2.**
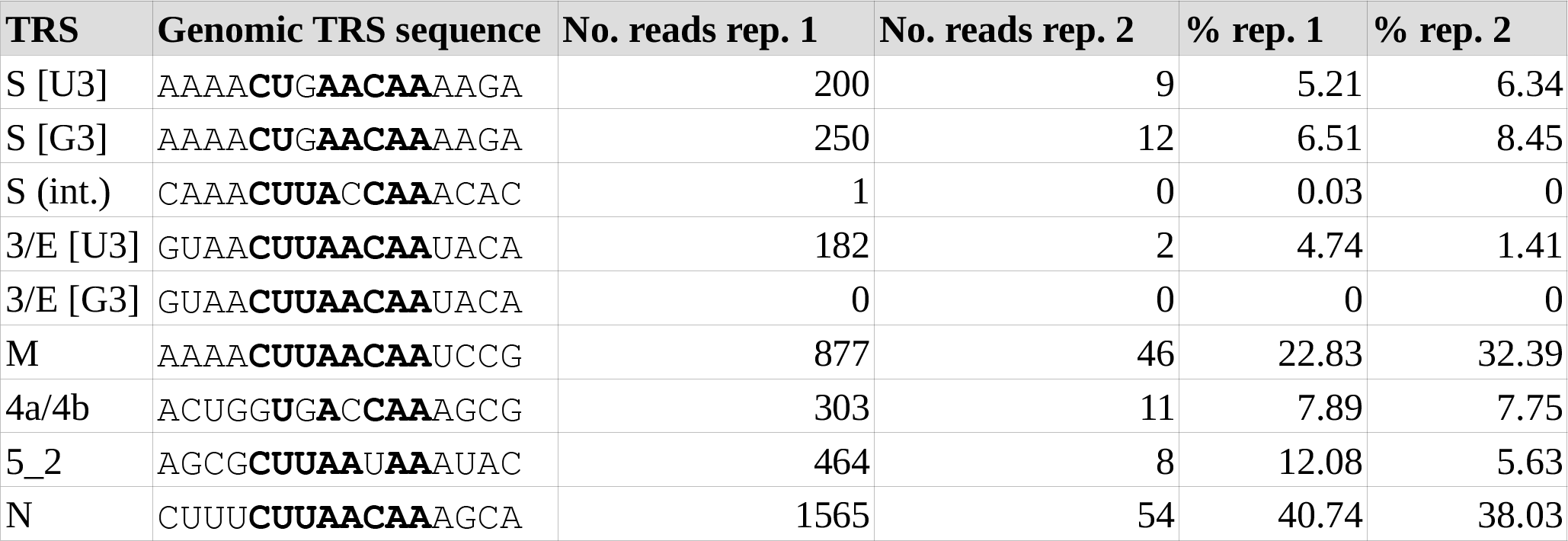
**(B)** Chimeric TRS-spanning reads identified in IBV M41 samples. Residues matching the leader TRS (TRS-L) are shown in bold.

**Table S3.** Top 20 most significantly enriched GO terms among genes which are down-regulated in IBV M41-infected cells relative to IBV Beaudette-infected cells. Column “Annotated” shows the total number of genes in the data set which are members of that GO term; “Significant” shows the number that are significantly differentially expressed; and “Expected” shows the proportion of genes that would be expected to occur in a random sample.

| GO.ID | Term | Annotated | Significant | Expected | P-value |
| --- | --- | --- | --- | --- | --- |
| GO:0010469 | regulation of signaling receptor activity | 53 | 7 | 0.58 | $1.2 \times 10^{-6}$ |
| GO:0048661 | positive regulation of smooth muscle cell proliferation | 18 | 5 | 0.2 | $3.5 \times 10^{-5}$ |
| GO:2000352 | negative regulation of endothelial cell apoptotic process | 9 | 3 | 0.1 | $9.7 \times 10^{-5}$ |
| GO:0060585 | positive regulation of prostaglandin-endoperoxide synthase | 2 | 2 | 0.02 | $1.2 \times 10^{-4}$ |
| GO:0033141 | positive regulation of peptidyl-serine phosphorylation of ST... | 2 | 2 | 0.02 | $1.2 \times 10^{-4}$ |
| GO:0014826 | vein smooth muscle contraction | 2 | 2 | 0.02 | $1.2 \times 10^{-4}$ |
| GO:1900625 | positive regulation of monocyte aggregation | 2 | 2 | 0.02 | $1.2 \times 10^{-4}$ |
| GO:0014824 | artery smooth muscle contraction | 3 | 2 | 0.03 | $3.5 \times 10^{-4}$ |
| GO:0003100 | regulation of systemic arterial blood pressure by endothelin | 3 | 2 | 0.03 | $3.5 \times 10^{-4}$ |
| GO:1901842 | negative regulation of high voltage-gated calcium channel ac... | 3 | 2 | 0.03 | $3.5 \times 10^{-4}$ |
| GO:0006955 | immune response | 244 | 12 | 2.66 | $5.9 \times 10^{-4}$ |
| GO:1904754 | positive regulation of vascular associated smooth muscle cel... | 4 | 2 | 0.04 | $6.9 \times 10^{-4}$ |
| GO:0071356 | cellular response to tumor necrosis factor | 40 | 4 | 0.44 | $8.4 \times 10^{-4}$ |
| GO:0043154 | negative regulation of cysteine-type endopeptidase activity ... | 18 | 3 | 0.2 | $8.8 \times 10^{-4}$ |
| GO:0001516 | prostaglandin biosynthetic process | 8 | 3 | 0.09 | $1.09 \times 10^{-3}$ |
| GO:0071498 | cellular response to fluid shear stress | 8 | 3 | 0.09 | $1.09 \times 10^{-3}$ |
| GO:0002931 | response to ischemia | 5 | 2 | 0.05 | $1.14 \times 10^{-3}$ |
| GO:0090051 | negative regulation of cell migration involved in sprouting ... | 5 | 2 | 0.05 | $1.14 \times 10^{-3}$ |
| GO:0019229 | regulation of vasoconstriction | 5 | 2 | 0.05 | $1.14 \times 10^{-3}$ |
| GO:0070373 | negative regulation of ERK1 and ERK2 cascade | 22 | 3 | 0.24 | $1.16 \times 10^{-3}$ |

**Table S4.** Top 20 most significantly enriched GO terms among genes which are up-regulated in IBV M41-infected cells relative to IBV Beaudette-infected cells. Column “Annotated” shows the total number of genes in the data set which are members of that GO term; “Significant” shows the number that are significantly differentially expressed; and “Expected” shows the proportion of genes that would be expected to occur in a random sample.

| GO.ID | Term | Annotated | Significant | Expected | P-value |
| --- | --- | --- | --- | --- | --- |
| GO:0042026 | protein refolding | 7 | 2 | 0.01 | $4.4 \times 10^{-5}$ |
| GO:0051085 | chaperone cofactor-dependent protein refolding | 9 | 2 | 0.01 | $7.6 \times 10^{-5}$ |
| GO:0034605 | cellular response to heat | 17 | 2 | 0.03 | $2.9 \times 10^{-4}$ |
| GO:0090074 | negative regulation of protein homodimerization activity | 1 | 1 | 0 | $1.6 \times 10^{-3}$ |
| GO:1905719 | protein localization to perinuclear region of cytoplasm | 1 | 1 | 0 | $1.6 \times 10^{-3}$ |
| GO:1903895 | negative regulation of IRE1-mediated unfolded protein respon... | 1 | 1 | 0 | $1.6 \times 10^{-3}$ |
| GO:0070194 | synaptonemal complex disassembly | 1 | 1 | 0 | $1.6 \times 10^{-3}$ |
| GO:0006880 | intracellular sequestering of iron ion | 1 | 1 | 0 | $1.6 \times 10^{-3}$ |
| GO:0034620 | cellular response to unfolded protein | 36 | 3 | 0.06 | $2.5 \times 10^{-3}$ |
| GO:1902949 | positive regulation of tau-protein kinase activity | 2 | 1 | 0 | $3.2 \times 10^{-3}$ |
| GO:1903071 | positive regulation of ER-associated ubiquitin-dependent pro... | 2 | 1 | 0 | $3.19 \times 10^{-3}$ |
| GO:1901896 | positive regulation of calcium-transporting ATPase activity | 2 | 1 | 0 | $3.19 \times 10^{-3}$ |
| GO:0090084 | negative regulation of inclusion body assembly | 3 | 1 | 0 | $4.78 \times 10^{-3}$ |
| GO:1905323 | telomerase holoenzyme complex assembly | 3 | 1 | 0 | $4.78 \times 10^{-3}$ |
| GO:0044130 | negative regulation of growth of symbiont in host | 3 | 1 | 0 | $4.78 \times 10^{-3}$ |
| GO:0007141 | male meiosis I | 3 | 1 | 0 | $4.78 \times 10^{-3}$ |
| GO:0031396 | regulation of protein ubiquitination | 73 | 2 | 0.12 | $5.31 \times 10^{-3}$ |
| GO:0035973 | aggrephagy | 4 | 1 | 0.01 | $6.38 \times 10^{-3}$ |
| GO:0061635 | regulation of protein complex stability | 4 | 1 | 0.01 | $6.38 \times 10^{-3}$ |
| GO:1900273 | positive regulation of long-term synaptic potentiation | 4 | 1 | 0.01 | $6.38 \times 10^{-3}$ |

**Figure S1.**
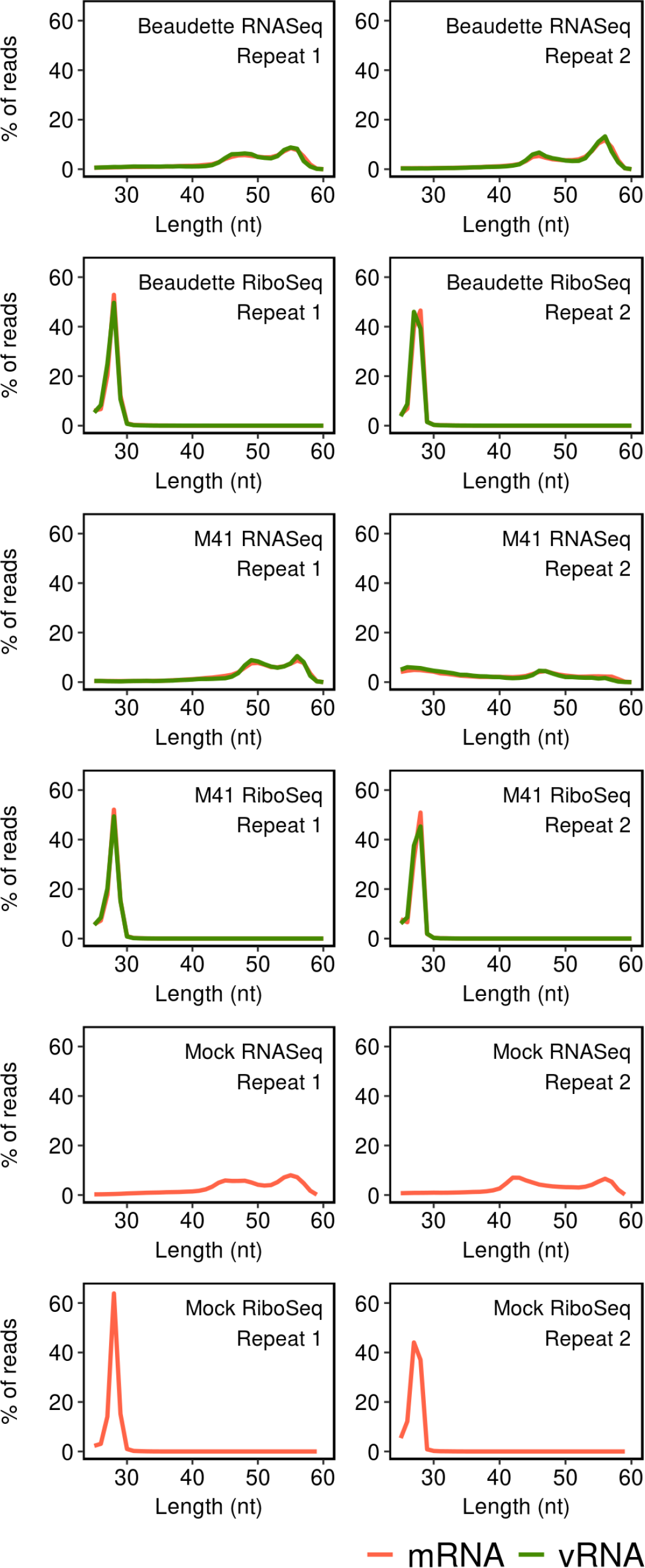
Length distribution of reads mapped to the internal regions of protein-coding sequences on viral RNA (vRNA; green lines) and host mRNA (red lines).

**Figure S2.**
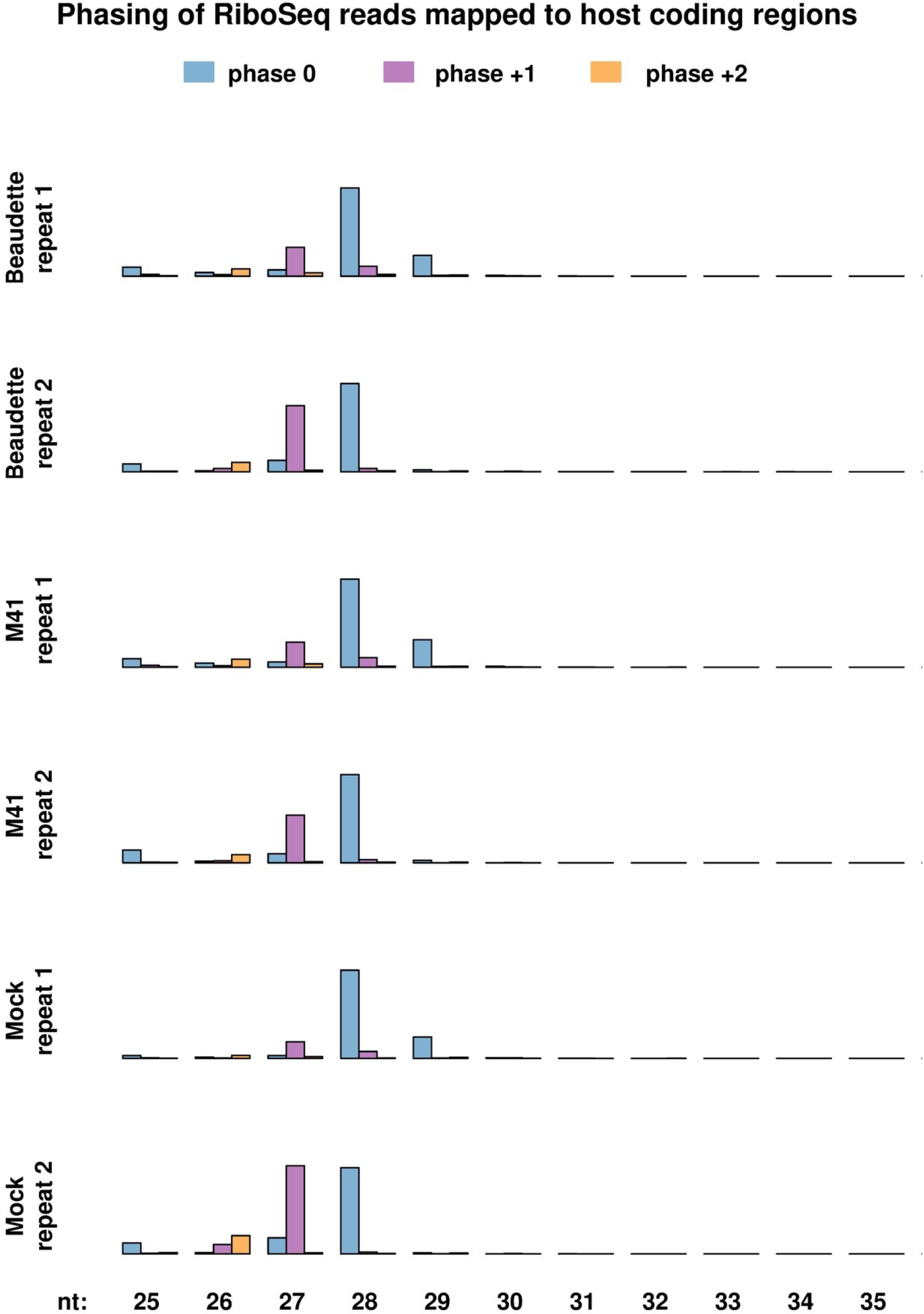
Phasing of RiboSeq reads mapped to the internal regions of protein-coding sequences in host mRNAs for different RPF lengths.

**Figure S3.**
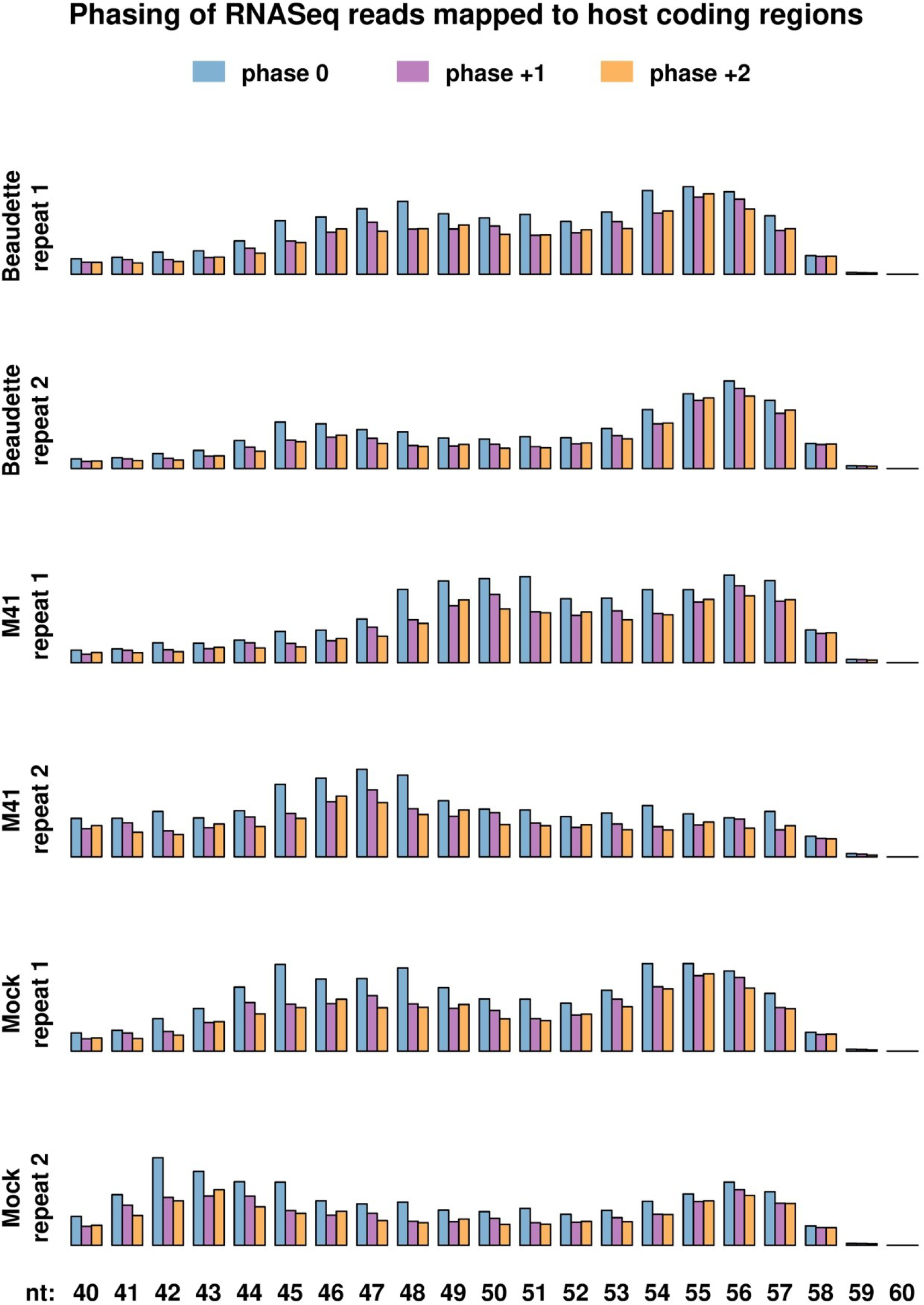
Phasing of RNASeq reads mapped to the internal regions of protein coding sequences in host mRNAs. To aid visualisation, only reads between 40 nt and 60 nt in length are shown.

**Figure S4.**
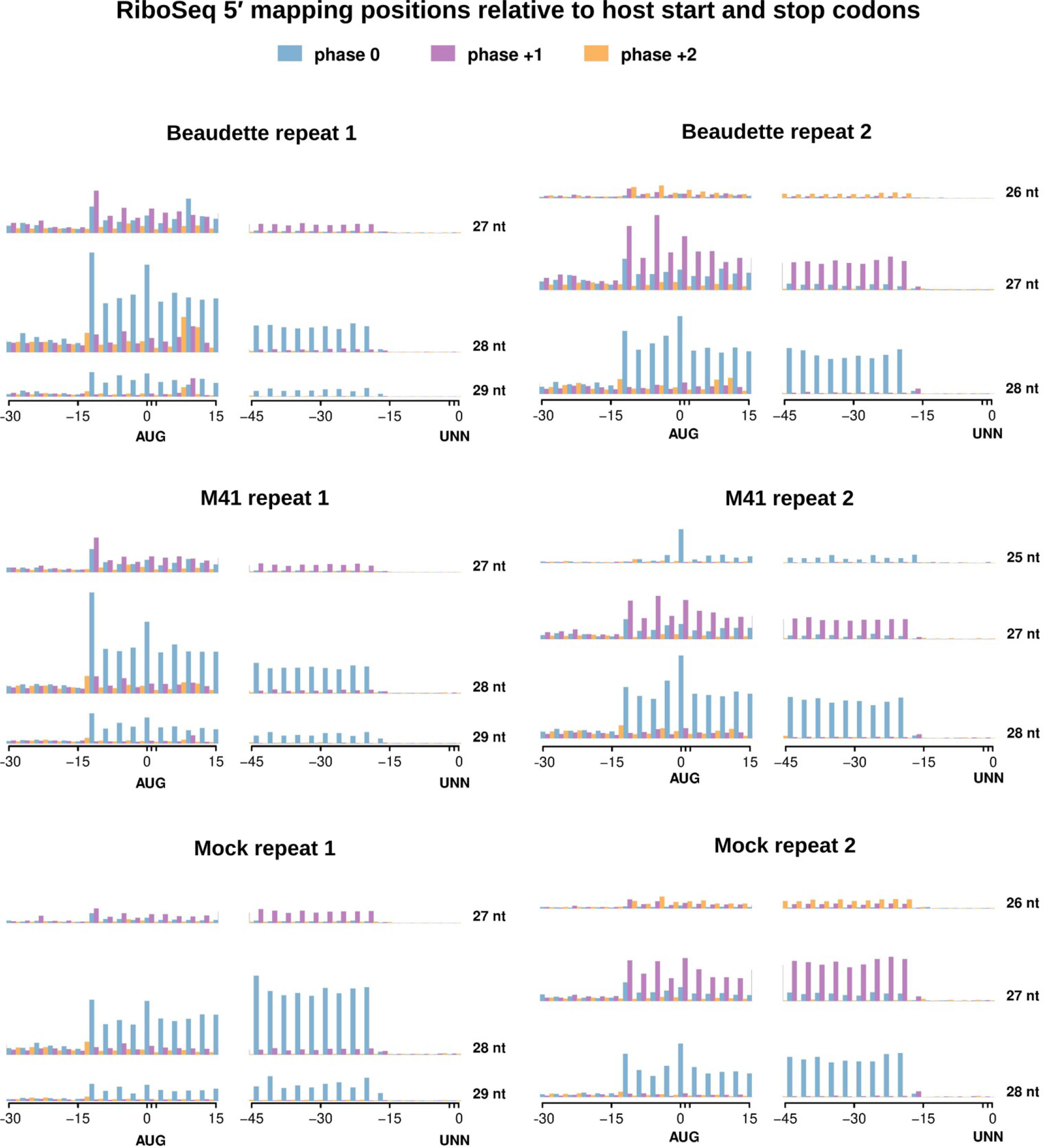
Histograms of 5′ mapping positions of RiboSeq reads relative to host mRNA start (AUG) and stop (UNN) codons. Positions indicated are relative to the first nt of the AUG codon (left) and the last nt of the UNN codon (right). The three most abundant read lengths are plotted for each library.

**Figure S5.**
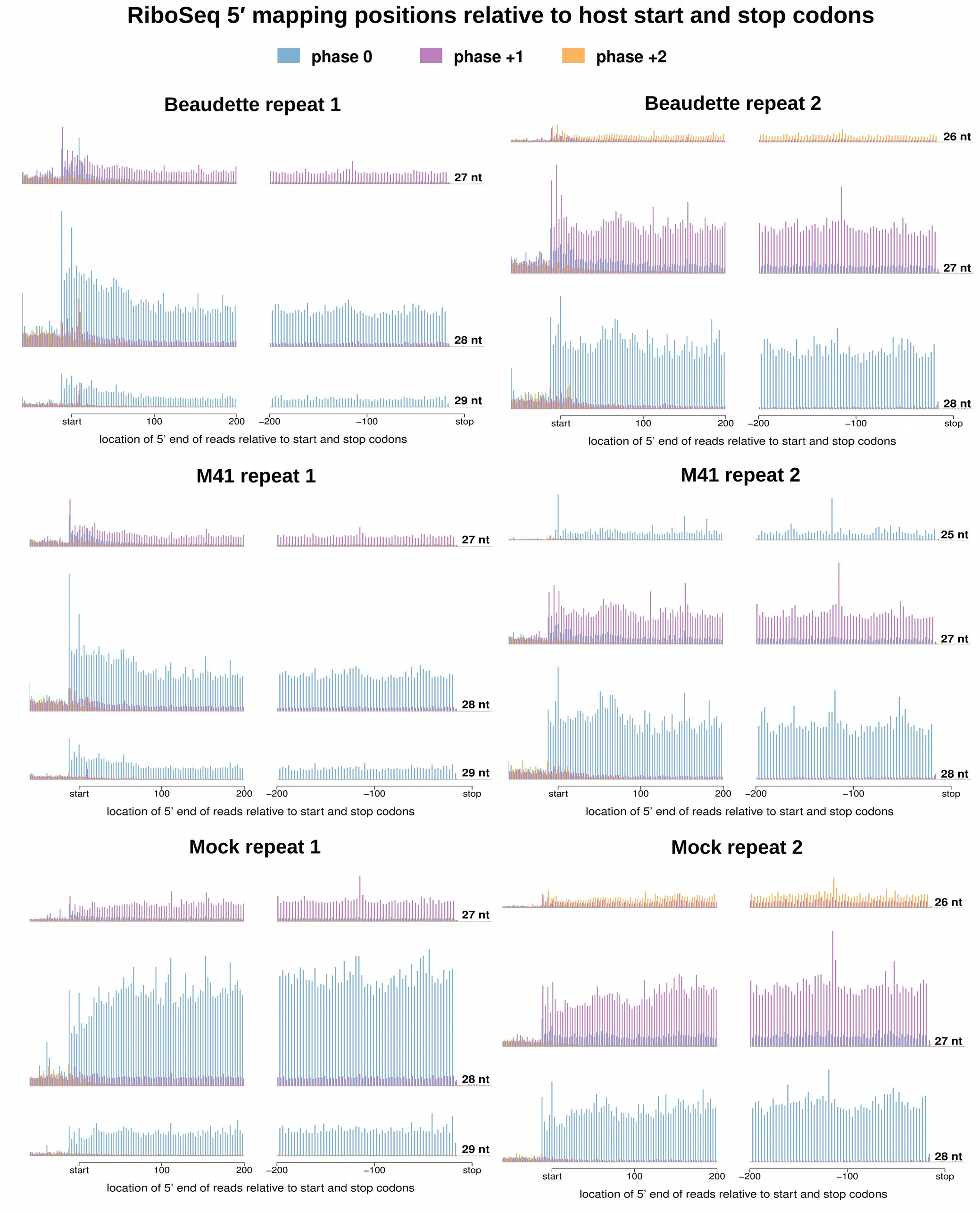
Histograms of 5′ mapping positions of RiboSeq reads relative to host mRNA start and stop codons. The three most abundant read lengths are plotted for each library.

**Figure S6.**
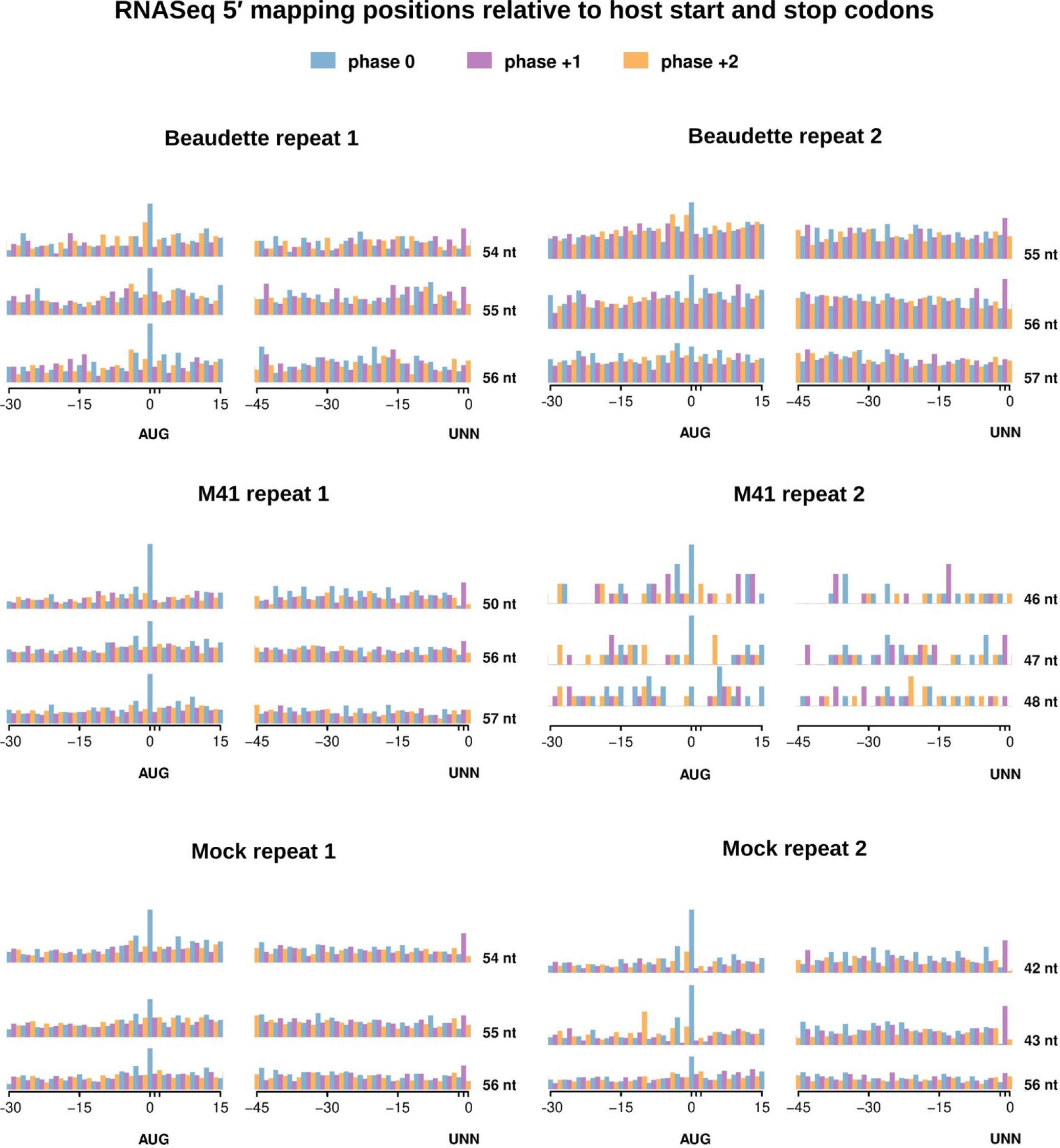
Histograms of 5′ mapping positions of RNASeq reads relative to host mRNA start (AUG) and stop (UNN) codons. Positions indicated are relative to the first nt of the AUG codon (left) and the last nt of the UNN codon (right). The three most abundant read lengths are plotted for each library.

**Figure S7.**
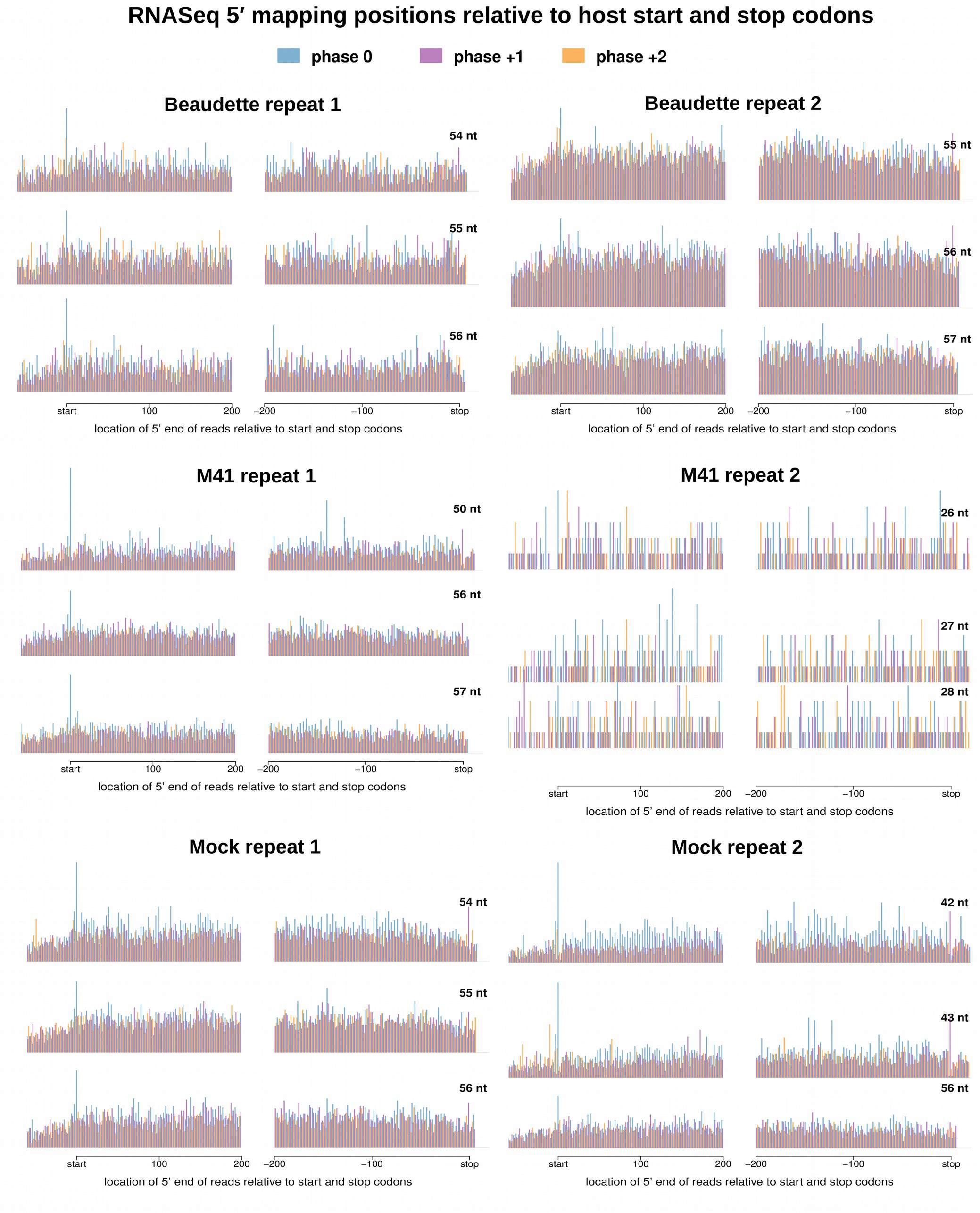
Histograms of 5′ mapping positions of RNASeq reads relative to host mRNA start and stop codons. The three most abundant read lengths are plotted for each library.

**Figure S8.**
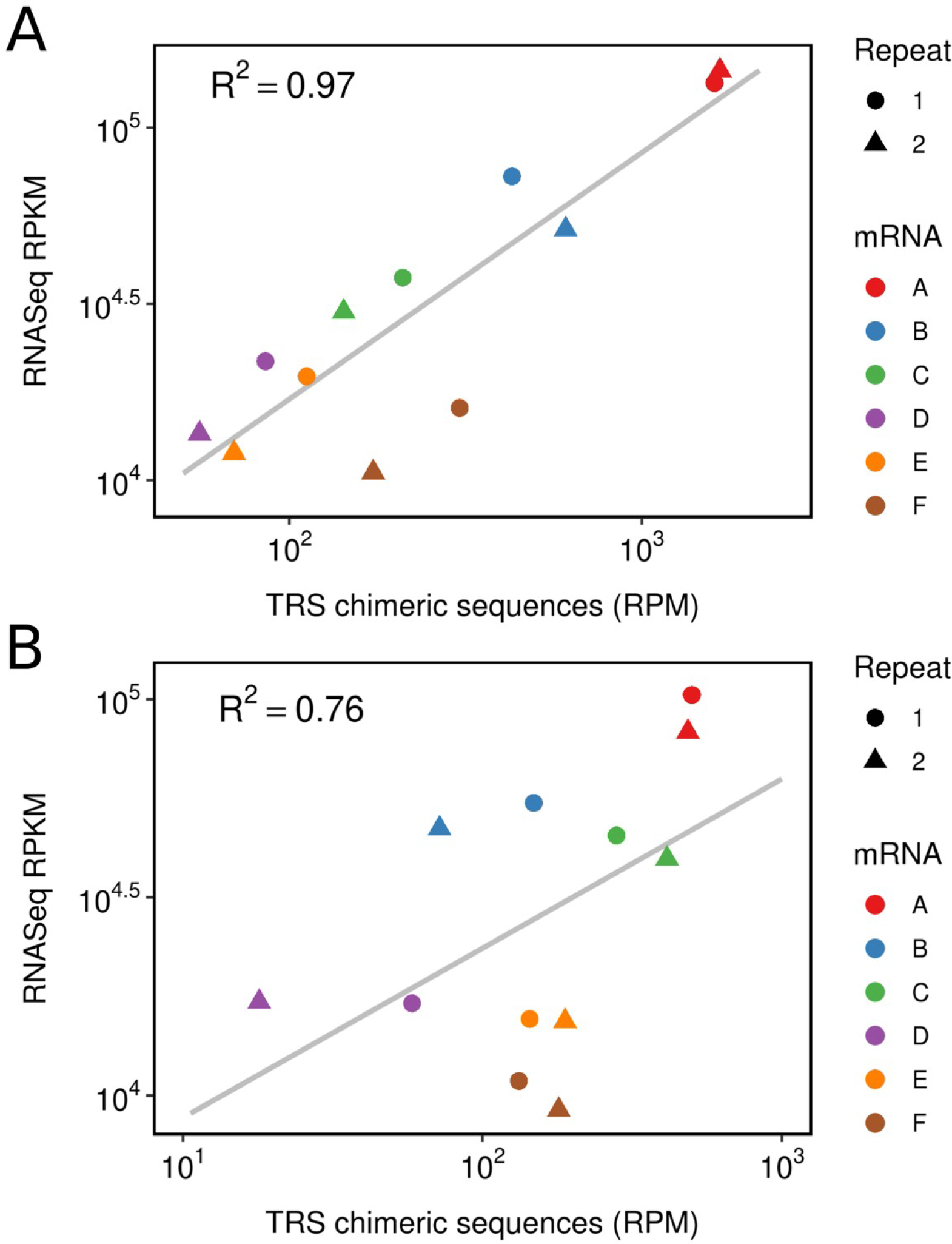
Relative abundances of sgmRNAs (mRNA A to E) and gRNA (mRNA F) for **(A)** IBV Beaudette and **(B)** IBV M41, as measured by decumulating RNASeq coverage (see Methods) or by counting chimeric reads spanning the TRS sequence. RNASeq densities are expressed as reads per kilobase per million mapped reads (RPKM), and chimeric TRS-spanning reads are expressed as reads per million mapped reads (RPM).

**Figure S9.**
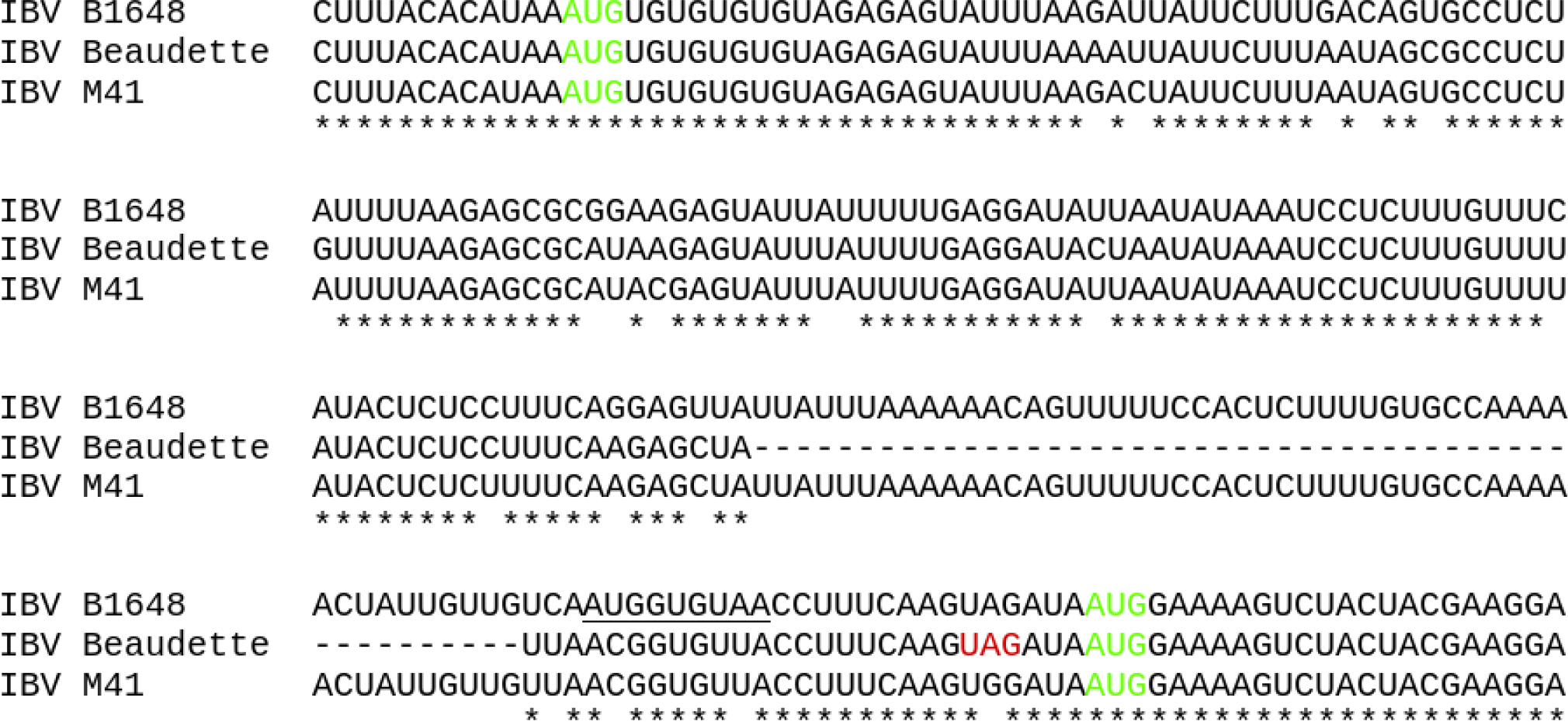
Alignment of IBV sequences, showing the positions of the ORF4b and ORF4c initiation codons (green). A 49-nt deletion in the Beaudette strain prematurely truncates the ORF4b gene by bringing a UAG codon (red) in frame. Note the absence of AUG codons between the starts of ORF4b and ORF4c in both IBV Beaudette and IBV M41, consistent with a leaky scanning mechanism for 4c expression. Conversely, the genome of the Belgian nephropathogenic strain B1648 contains an intervening AUG (underlined; 2-codon ORF) towards the 3′ end of ORF4b.

**Figure S10.**
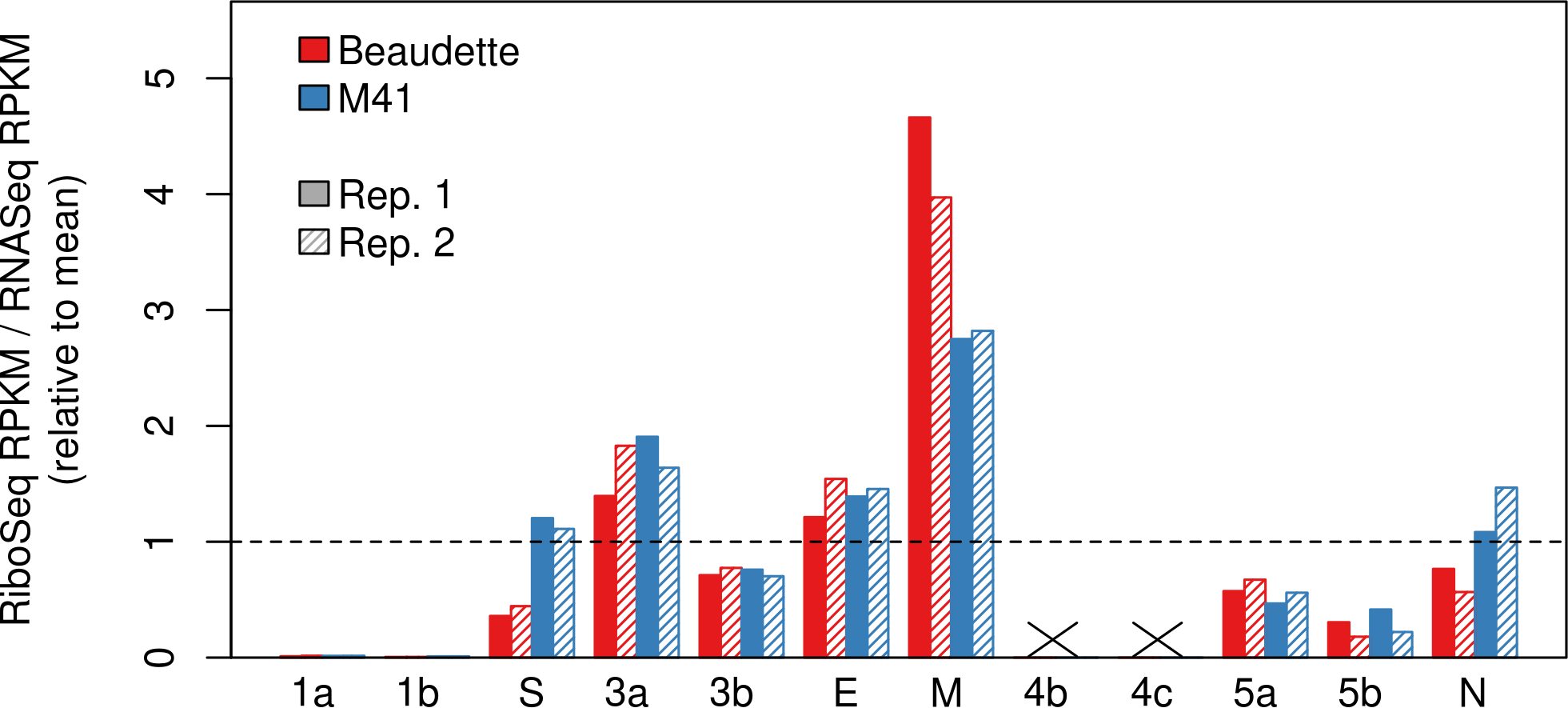
Viral gene translation efficiency values calculated using decumulated RNASeq reads (expressed as reads per kilobase per million mapped reads [RPKM]). The ratio of RiboSeq RPKM to RNASeq RPKM is plotted relative to the mean across all samples.

**Figure S11.**
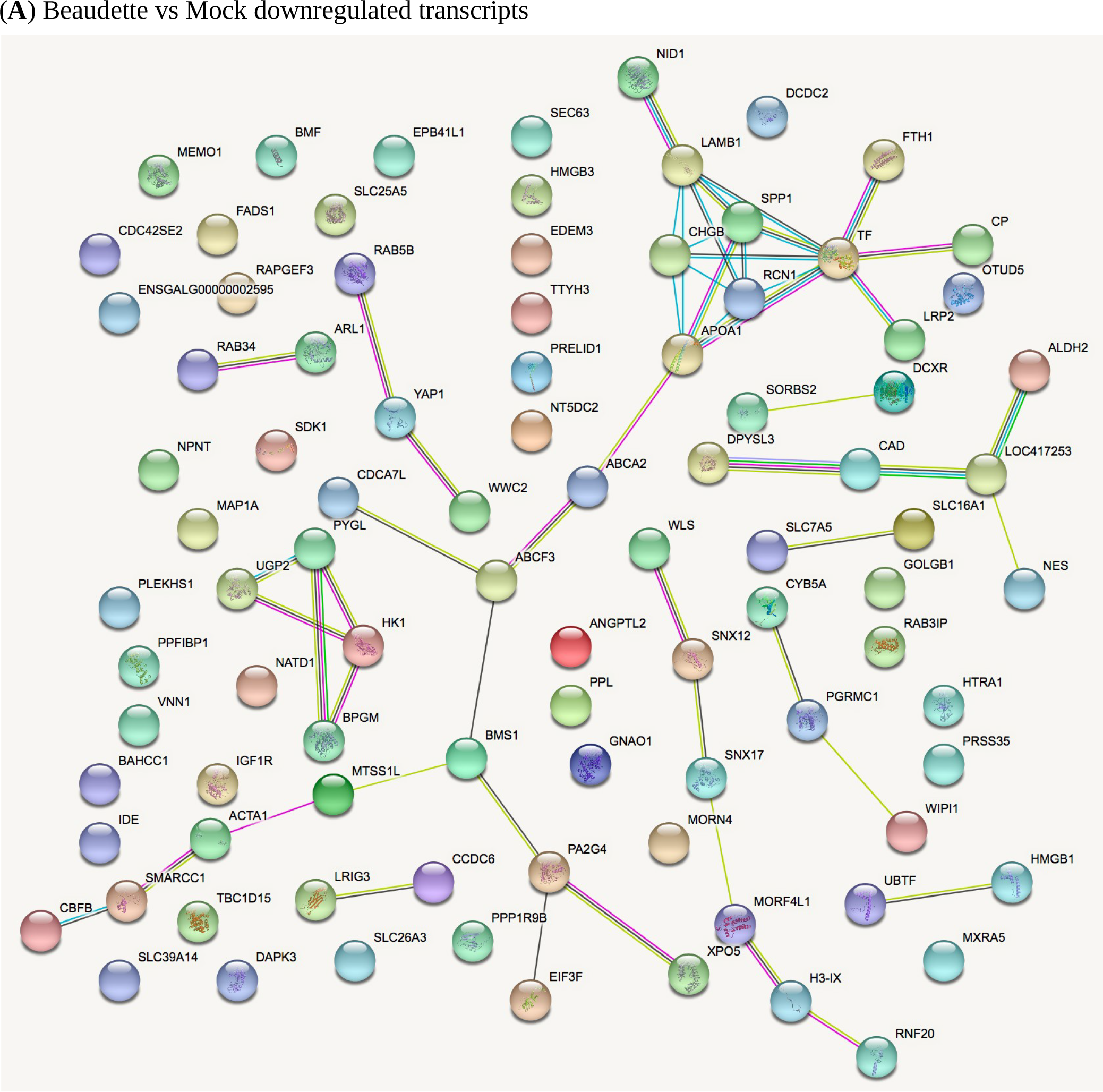
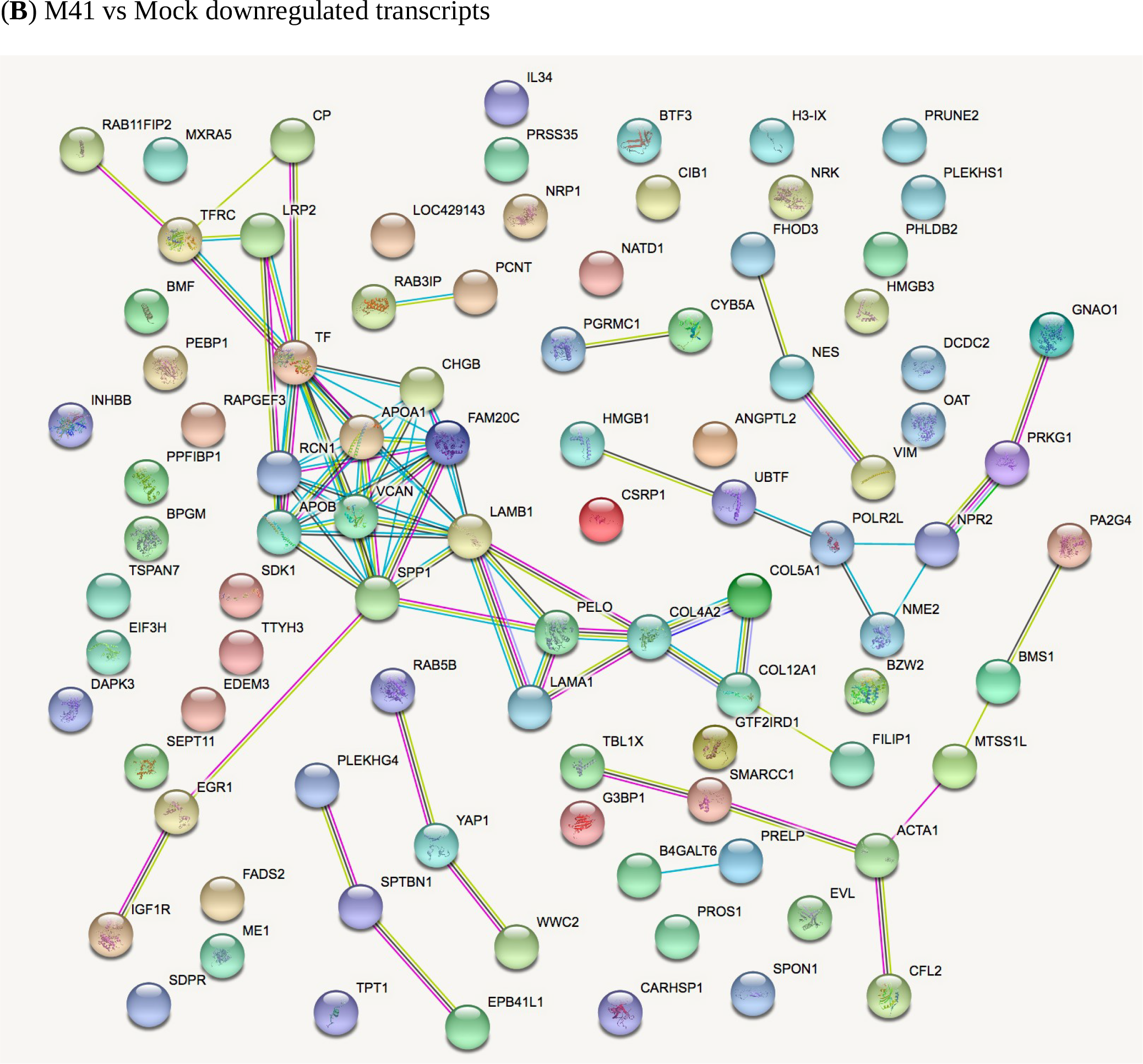
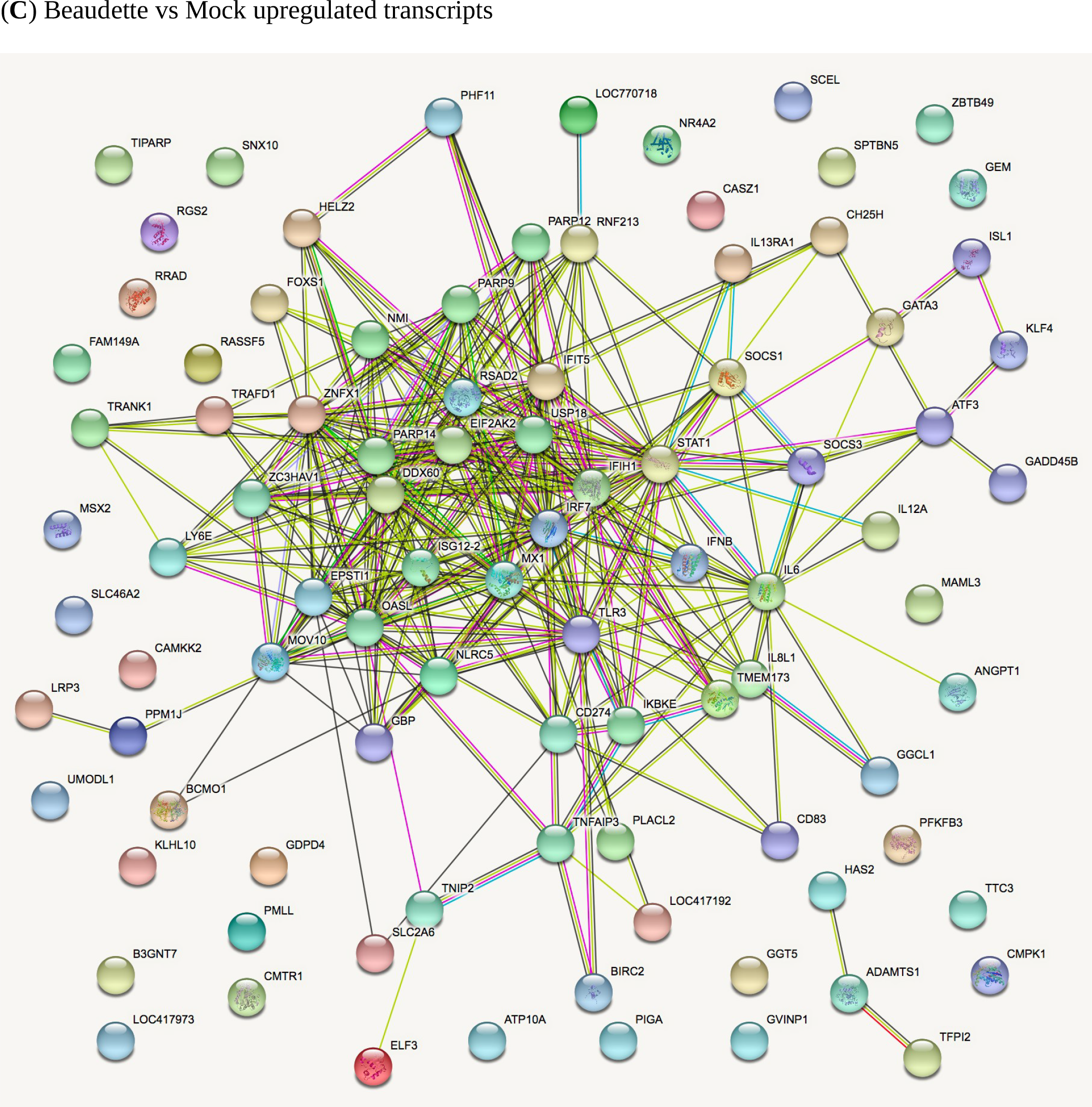
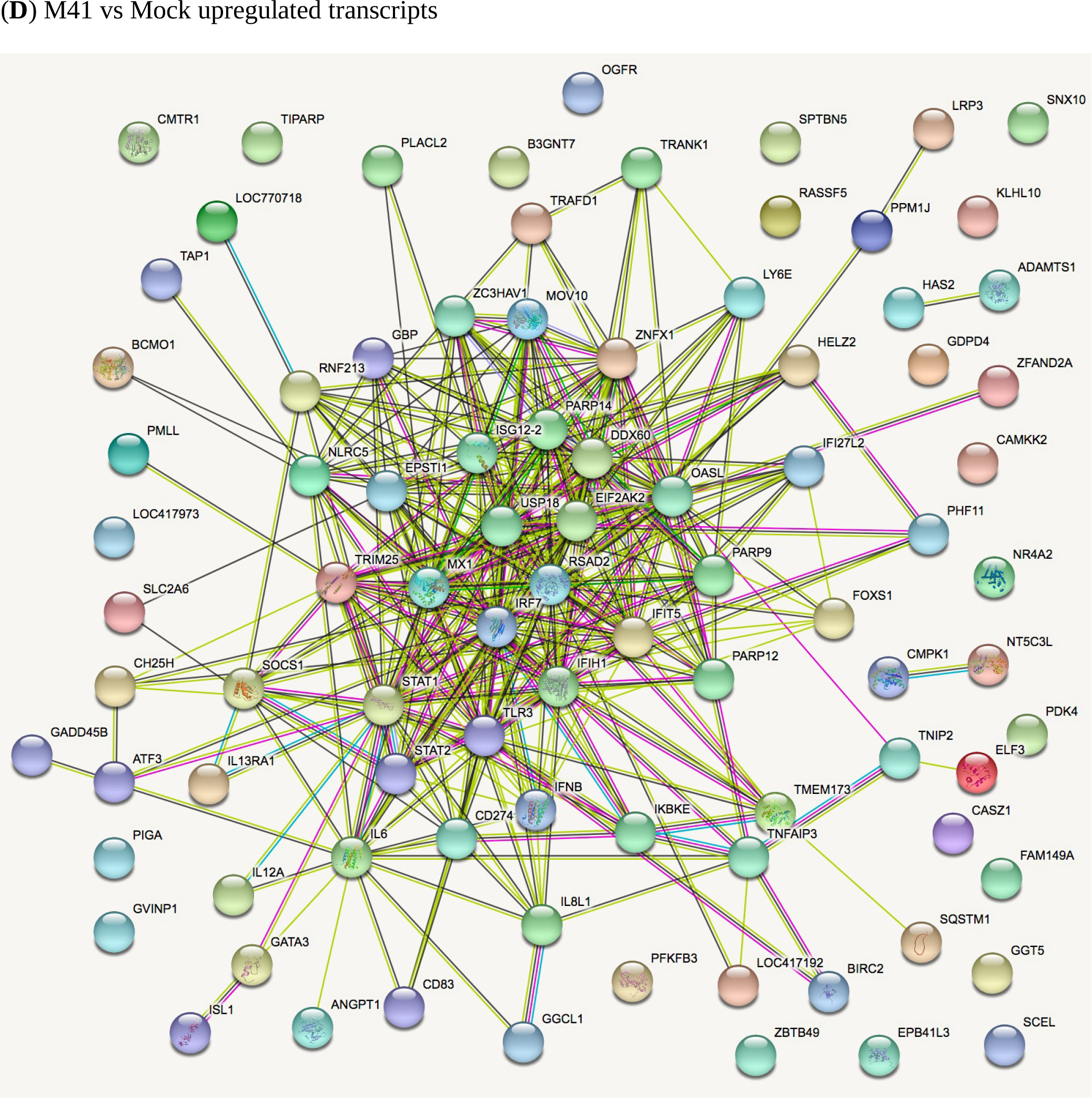
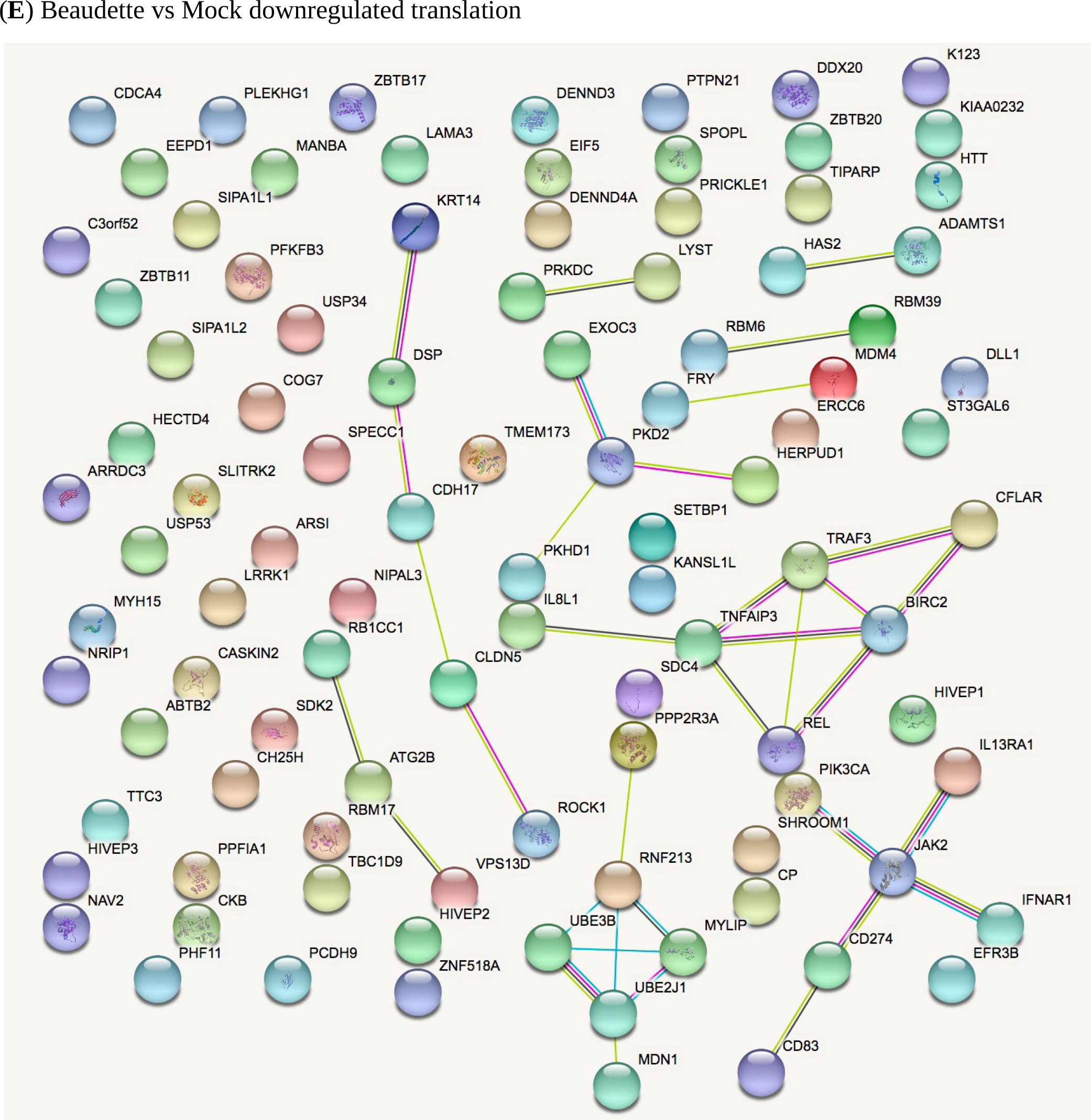
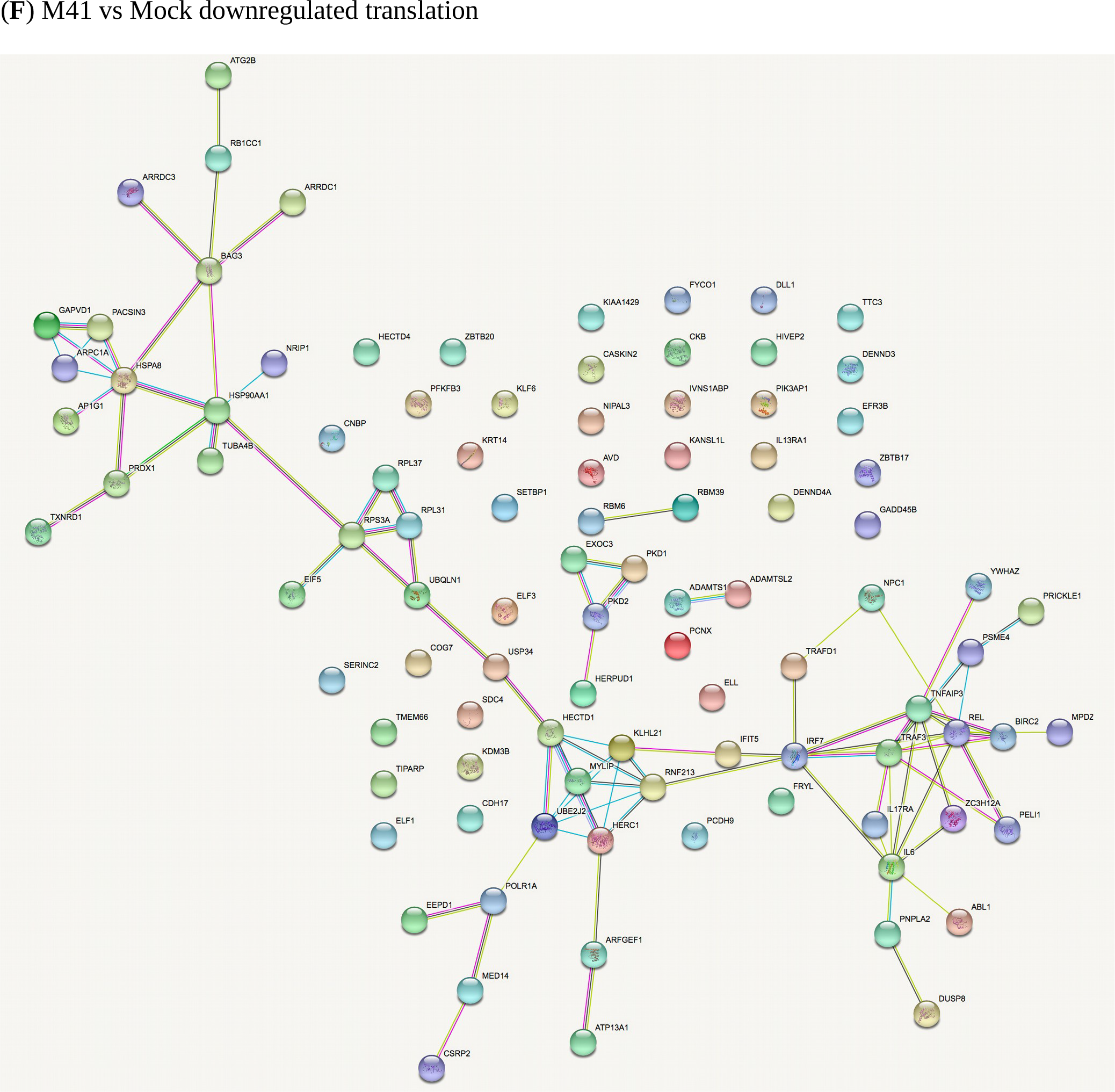
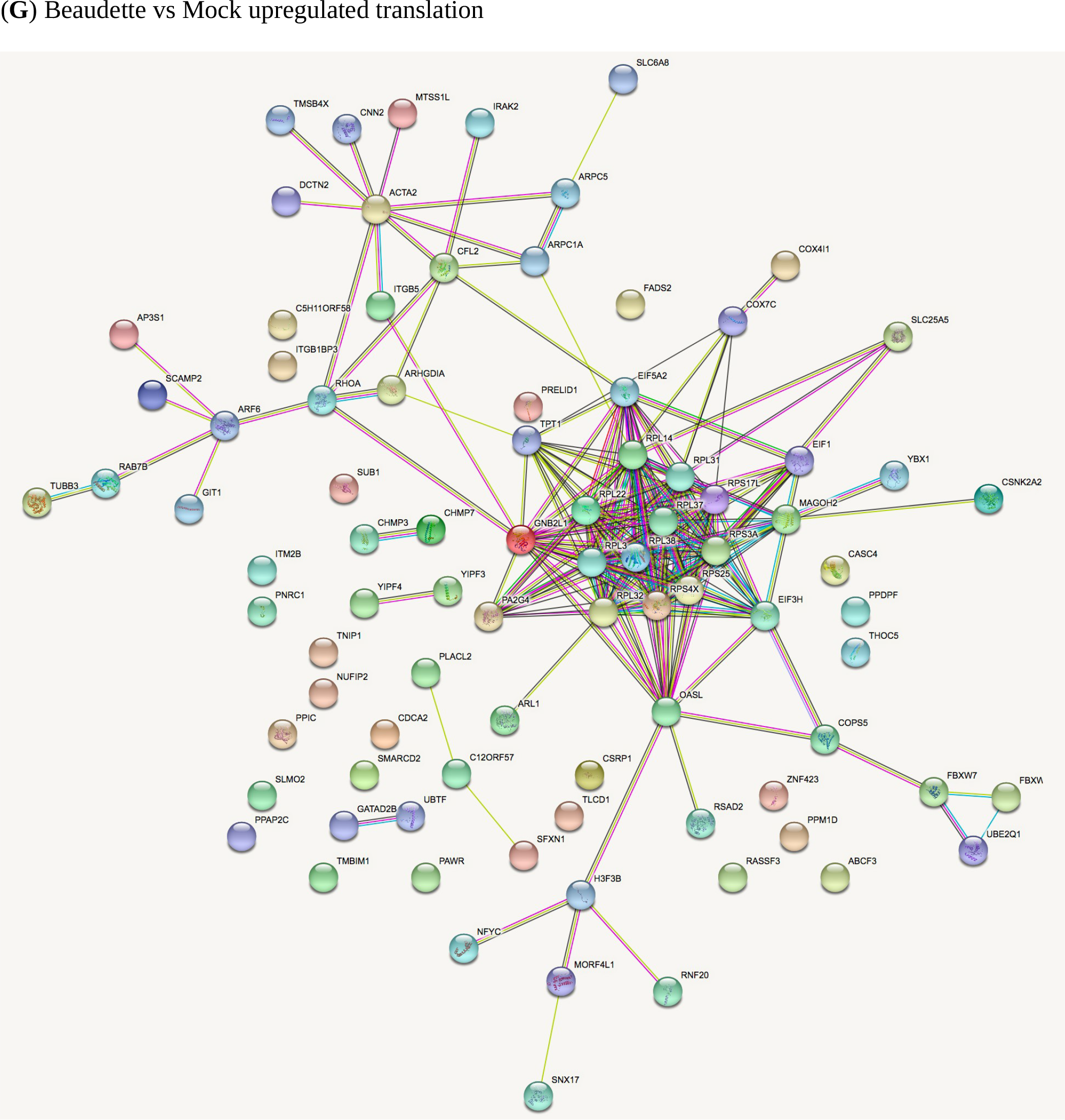
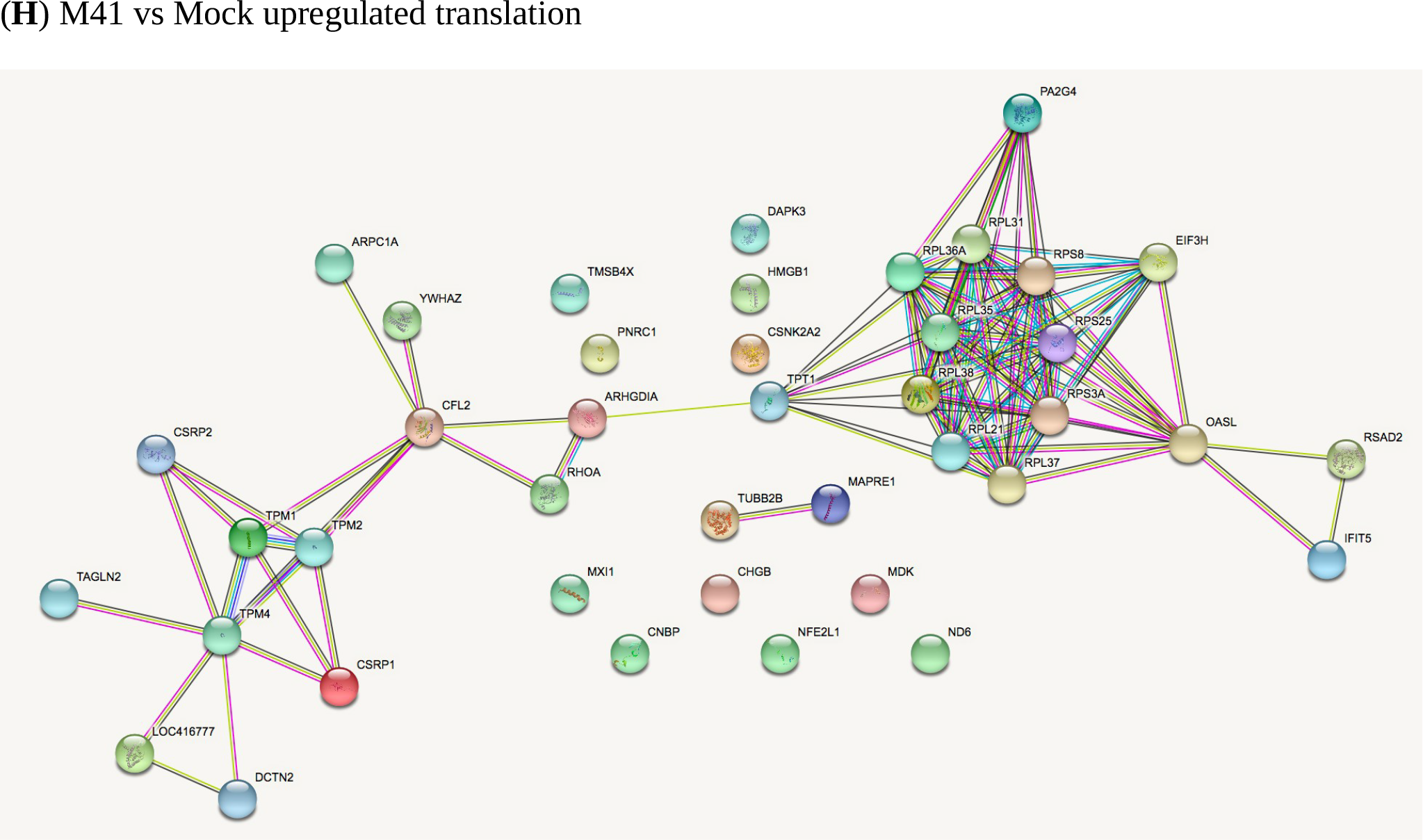
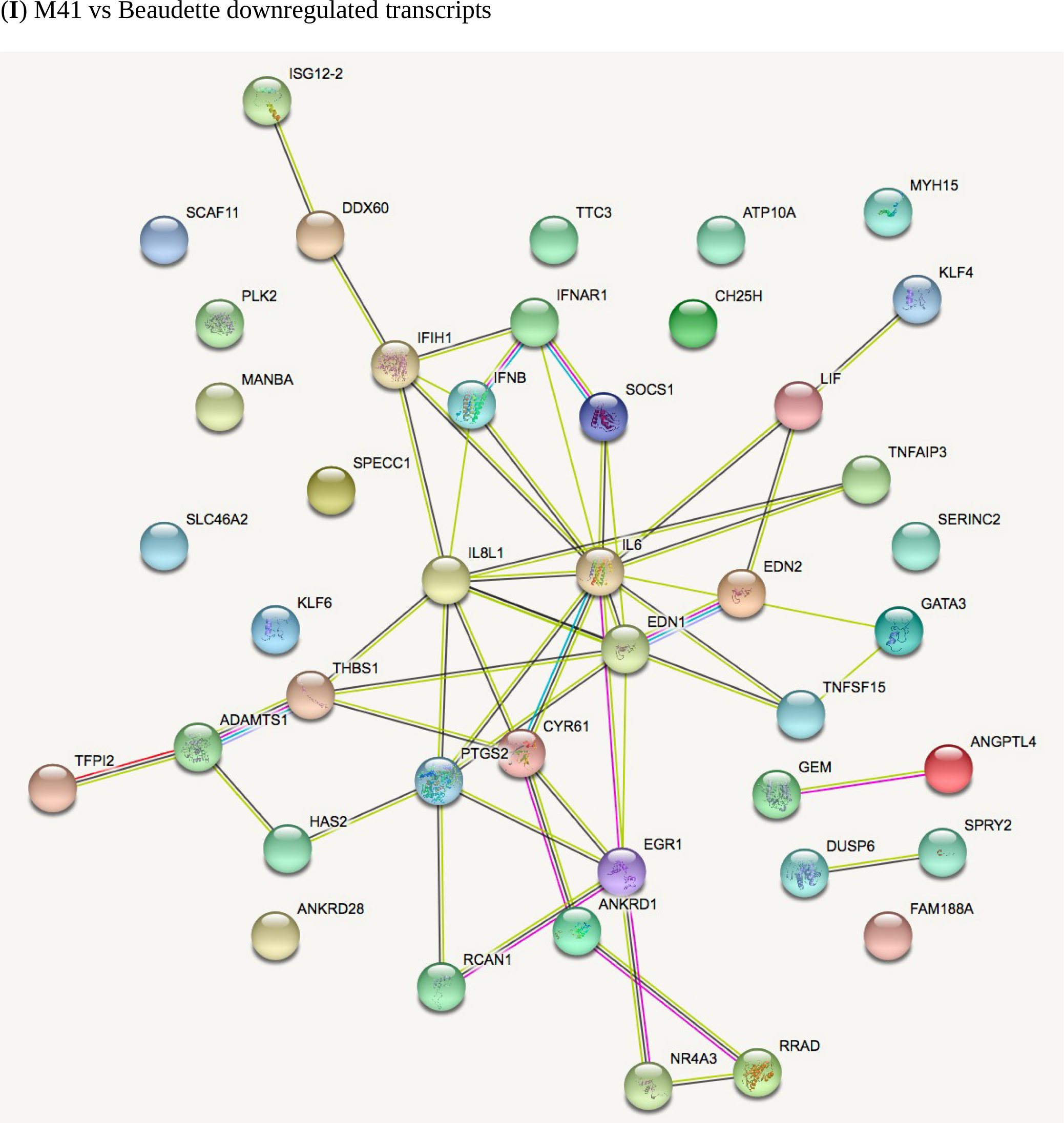
STRING analysis of the relationship between differentially expressed genes in comparisons of IBV Beaudette, IBV M41 and mock-infected chick kidney cells (**panels A to I**). The network nodes represent the proteins encoded by the differentially expressed genes. Seven different coloured lines link a number of nodes and represent seven types of evidence used in predicting associations. A red line indicates the presence of fusion evidence; a green line represents neighborhood evidence; a blue line represents co-ocurrence evidence; a purple line represents experimental evidence; a yellow line represents text-mining evidence; a light blue line represents database evidence; and a black line represents co-expression evidence.

## References

Aird, D., Ross, M.G., Chen, W.S., Danielsson, M., Fennell, T., Russ, C., Jaffe, D.B., Nusbaum, C. and Gnirke, A. (2011). Analyzing and minimizing PCR amplification bias in Illumina sequencing libraries. Genome Biol. 12: R18.

Alexa A, Rahnenfuhrer J (2018). topGO: Enrichment Analysis for Gene Ontology. R package version 2.34.0.

Alonso-Caplen, F.V., Matsuoka, Y., Wilcox, G.E. and Compans, R.W. (1984). Replication and morphogenesis of avian coronavirus in Vero cells and their inhibition by monensin. Virus Res. 1:153–167.

An, H., Cai, Z., Yang, Y., Wang, Z., Liu, D.X. and Fang, S. (2019). Identification and formation mechanism of a novel noncoding RNA produced by avian infectious bronchitis virus. Virology 528:176–180.

Anders, S., Pyl, P.T. and Huber, W. (2015). HTSeq - a Python framework to work with high-throughput sequencing data. Bioinformatics 31:166–169.

Atkins, J.F., Loughran, G., Bhatt, P.R., Firth, A.E. and Baranov, P.V. (2016). Ribosomal frameshifting and transcriptional slippage: From genetic steganography and cryptography to adventitious use. Nucleic Acids Res. 44:7007–7078.

Beaudette, F. R. and Hudson C. R. (1937). Cultivation of the virus of infectious bronchitis. J. Am. Vet. Med. Assoc. 90: 51–60.

Bentley, K., Keep, S.M., Armesto, M. and Britton, P. (2013). Identification of a noncanonically transcribed subgenomic mRNA of infectious bronchitis virus and other gammacoronaviruses. J. Virol. 87: 2128–2136.

Bickerton, E., Maier, HJ., Stevenson-Leggett, P., Armesto, M. and Britton, P. (2018). The S2 Subunit of Infectious Bronchitis Virus Beaudette Is a Determinant of Cellular Tropism. J. Virol. 92. pii: e01044–18.

Boursnell, M.E., Brown, T.D., Foulds, I.J., Green, P.F., Tomley, F.M., Binns, M.M. (1987). Completion of the sequence of the genome of the coronavirus avian infectious bronchitis virus. J. Gen. Virol. 68:57–77.

Brierley, I., Boursnell, M.E., Binns, M.M., Bilimoria, B., Blok, V.C., Brown, T.D. and Inglis, S.C. (1987). An efficient ribosomal frame-shifting signal in the polymerase-encoding region of the coronavirus IBV. EMBO J. 6: 3779–3785.

Brierley, I, Digard, P. and Inglis, S.C. (1989). Characterization of an efficient coronavirus ribosomal frameshifting signal: requirement for an RNA pseudoknot. Cell 57: 537–547.

Brierley I. and Pennell, S. (2001). Structure and function of the stimulatory RNAs involved in programmed eukaryotic-1 ribosomal frameshifting. Cold Spring Harb. Symp. Quant. Biol. 66: 233–248.

Brockway, S.M, Clay, C.T., Lu, X.T. and Denison, M.R. (2003). Characterization of the expression, intracellular localization, and replication complex association of the putative mouse hepatitis virus RNA-dependent RNA polymerase. J. Virol. 77: 10515–10527.

Brown, T.D., Boursnell, M.E., Binns, M.M. and Tomley, F.M. (1986). Cloning and sequencing of 5’ terminal sequences from avian infectious bronchitis virus genomic RNA. J. Gen. Virol. 67:221–228.

Cao, J., Wu, C.C. and Lin, T.L. (2008). Complete nucleotide sequence of polyprotein gene 1 and genome organization of turkey coronavirus. Virus Res. 136: 43–49.

Casais, R., Dove, B., Cavanagh D. and Britton, P. (2003). Recombinant avian infectious bronchitis virus expressing a heterologous spike gene demonstrates that the spike protein is a determinant of cell tropism. J. Virol. 77: 9084–9089.

Casais, R., Thiel, V., Siddell, S.G., Cavanagh, D. and Britton, P. (2001). Reverse genetics system for the avian coronavirus infectious bronchitis virus. J. Virol. 75, 12359–12369.

Cavanagh, D. (2005). Coronaviruses in poultry and other birds. Avian Pathol. 34: 439–448.

Cavanagh, D. (2007). Coronavirus avian infectious bronchitis virus. Vet. Res. 38: 281–297.

Chadani, Y., Niwa, T., Chiba, S., Taguchi, H. and Ito, K. (2016). Integrated *in vivo* and *in vitro* nascent chain profiling reveals widespread translational pausing. Proc. Natl. Acad. Sci. U.S.A. 113:E829–38.

Chhabra, R., Ball, C., Chantrey, J. and Ganapathy, K. (2018). Differential innate immune responses induced by classical and variant infectious bronchitis viruses in specific pathogen free chicks. Dev. Comp. Immunol. 87:16–23.

Charneski, C.A. and Hurst, L.D. (2013). Positively charged residues are the major determinants of ribosomal velocity. PLoS Biol. 11:e1001508.

Chung, B.Y., Hardcastle, T.J., Jones J..D, Irigoyen,. N, Firth, A.E., Baulcombe, D.C. and Brierley, I. (2015). The use of duplex-specific nuclease in ribosome profiling and a user-friendly software package for Ribo-seq data analysis. RNA 21:1731–1745.

Cong, F., Liu, X., Han Z., Shao, Y., Kong, X. and Liu, S. (2013). Transcriptome analysis of chicken kidney tissues following coronavirus avian infectious bronchitis virus infection. BMC Genomics 14:743.

Cunningham, C.H., Spring, M.P. and Nazerian, K. (1972). Replication of avian infectious bronchitis virus in African green monkey kidney cell line VERO. J. Gen. Virol. 16: 423–427.

Cunningham, F., Achuthan, P., Akanni, W., Allen, J., Amode, M.R., Armean, I.M., Bennett, R., Bhai, J., Billis, K., Boddu, S. et al. (2019). Ensembl 2019. Nucleic Acids Res. 47:D745–D751.

Dobin, A., Davis, C.A., Schlesinger, F., Drenkow, J., Zaleski, C., Jha, S., Batut, P, Chaisson, M. and Gingeras, T.R. (2013). STAR: ultrafast universal RNA-seq aligner. Bioinformatics 29:15–21.

Doyle, N., Neuman, B.W., Simpson, J., Hawes, P.C., Mantell, J., Verkade, P., Alrashedi, H. and Maier, H.J. (2018). Infectious Bronchitis Virus Nonstructural Protein 4 Alone Induces Membrane Pairing. Viruses 10 pii: E477.

Duncan, C.D.S. and Mata, J. (2017). Effects of cycloheximide on the interpretation of ribosome profiling experiments in Schizosaccharomyces pombe. Sci. Rep. 7:10331.

Firth, A.E. and Brierley, I. (2012). Non-canonical translation in RNA viruses. J. Gen. Virol. 93:1385–1409.

Fuchs, R.T., Sun, Z., Zhuang, F. and Robb, G.B (2015). Bias in ligation-based small RNA sequencing library construction is determined by adaptor and RNA structure. PLoS One. 10: e0126049.

Gerashchenko, M.V. and Gladyshev, V.N. (2014). Translation inhibitors cause abnormalities in ribosome profiling experiments. Nucleic Acids Res. 42:e134.

Gomaa, M.H., Barta, J.R., Ojkic, D. and Yoo, D. (2008). Complete genomic sequence of turkey coronavirus. Virus Res. 135: 237–246.

Gorbalenya, A.E, Enjuanes, L., Ziebuhr, J. and Snijder E.J. (2006). Nidovirales: evolving the largest RNA virus genome. Virus Res. 117: 17–37.

Hennion, R. and Hill, G. (2015). In Coronaviruses Vol. 1282 Methods in Molecular Biology (eds Helena Jane Maier, Erica Bickerton, & Paul Britton) Ch. 6, 57–62. (Springer New York).

Hodgson, T., Casais, R., Dove, B., Britton, P. and Cavanagh, D. (2014). Recombinant infectious bronchitis coronavirus Beaudette with the spike protein gene of the pathogenic M41 strain remains attenuated but induces protective immunity. J. Virol. 78: 13804–13811.

Ingolia, N.T., Ghaemmaghami, S., Newman, J.R. and Weissman, J.S. (2009). Genome-wide analysis in vivo of translation with nucleotide resolution using ribosome profiling. Science. 324: 218–223.

Ingolia, N.T., Lareau, L.F. and Weissman, J.S. (2011). Ribosome profiling of mouse embryonic stem cells reveals the complexity and dynamics of mammalian proteomes. Cell. 147: 789–802.

Ingolia, N.T., Brar, G.A., Rouskin, S., McGeachy, A.M. and Weissman, J.S. (2012). The ribosome profiling strategy for monitoring translation in vivo by deep sequencing of ribosome-protected mRNA fragments. Nat. Protoc. 7:1534–1550.

Ingolia, N.T. (2014). Ribosome profiling: new views of translation, from single codons to genome scale. Nat. Rev. Genet. 15:205–213.

Irigoyen, N., Firth, A.E., Jones, J.D., Chung, B.Y., Siddell, S.G and Brierley, I. (2016). High-resolution analysis of coronavirus gene expression by RNA sequencing and ribosome profiling. PLoS Pathog. 12:e1005473.

Irigoyen, N., Dinan, A.M., Brierley, I. and Firth, A.E. (2018). Ribosome profiling of the retrovirus murine leukemia virus. Retrovirology 15: 10.

Jukes, T.H. (1996). On the prevalence of certain codons (“RNY”) in genes for proteins. J. Mol. Evol. 42:3 77–381.

Kottier, S.A., Cavanagh, D. and Britton, P. (1995). Experimental evidence of recombination in coronavirus infectious bronchitis virus. Virology. 213:569–580.

Kumari, R., Michel, A.M. and Baranov, P.V. (2018). PausePred and Rfeet: webtools for inferring ribosome pauses and visualizing footprint density from ribosome profiling data. RNA 24:1297–1304.

Kuo, L. and Masters, P.S. (2013). Functional analysis of the murine coronavirus genomic RNA packaging signal. J. Virol. 87:5182–5192.

Langmead, B., Trapnell, C., Pop, M. and Salzberg, S.L. (2009). Ultrafast and memory-efficient alignment of short DNA sequences to the human genome. Genome Biol. 10: R25.

Laude, Hubert, and Paul S. Masters. “The coronavirus nucleocapsid protein.” The Coronaviridae. Springer US, 1995. 141–163.

Liu, D.X., Xu, H.Y. and Brown, T.D. (1997). Proteolytic processing of the coronavirus infectious bronchitis virus 1a polyprotein: identification of a 10-kilodalton polypeptide and determination of its cleavage sites. J Virol. 71:1814–1820.

Oostra, M., te Lintelo, E.G., Deijs, M., Verheije, M.H., Rottier, P.J. and de Haan, C.A. (2007). Localization and membrane topology of coronavirus nonstructural protein 4: involvement of the early secretory pathway in replication. J. Virol. 81:12323–12336.

Otsuki, K., Noro, K., Yamamoto, H. and Tsubokura, M. (1979). Studies on avian infectious bronchitis virus (IBV). II. Propagation of IBV in several cultured cells. Arch. Virol. 60: 115–122.

Pasternak, A.O., van den Born, E., Spaan, W.J. and Snijder, E.J. (2001). Sequence requirements for RNA strand transfer during nidovirus discontinuous subgenomic RNA synthesis. EMBO J. 20: 7220–7228.

Reddy, V.R., Theuns, S., Roukaerts, I.D., Zeller, M., Matthijnssens, J. and Nauwynck, H.J. (2015). Genetic Characterization of the Belgian Nephropathogenic Infectious Bronchitis Virus (NIBV) Reference Strain B1648. Viruses 7: 4488–4506.

Requião, R.D., de Souza, H.J., Rossetto, S., Domitrovic, T. and Palhano, F.L. (2016). Increased ribosome density associated to positively charged residues is evident in ribosome profiling experiments performed in the absence of translation inhibitors. RNA Biol. 13:561–568.

Sabi, R. and Tuller T. (2015). A comparative genomics study on the effect of individual amino acids on ribosome stalling. BMC Genomics 16 Suppl. 10:S5.

Sabi, R. and Tuller, T. (2017). Computational analysis of nascent peptides that induce ribosome stalling and their proteomic distribution in *Saccharomyces cerevisiae*. RNA 23:983–994.

Sawicki, S.G. and Sawicki, D.L. (1995). Coronaviruses use discontinuous extension for synthesis of subgenome-length negative strands. Adv. Exp. Med. Biol. 380: 499–506.

Sawicki, S.G. and Sawicki, D.L. (1998). A new model for coronavirus transcription. Adv. Exp. Med. Biol. 440: 215–219.

Sawicki, S.G., Sawicki, D.L. and Siddell, S.G. (2007). A contemporary view of coronavirus transcription. J. Virol. 81: 20–29.

Stewart, H., Brown, K., Dinan, A.M., Irigoyen, N., Snijder, E.J. and Firth, A.E. (2018). Transcriptional and translational landscape of equine torovirus. J. Virol. 92 pii: e00589–18.

Sola, I., Moreno, J.L., Zúñiga, S., Alonso, S. and Enjuanes, L. (2005). Role of nucleotides immediately flanking the transcription-regulating sequence core in coronavirus subgenomic mRNA synthesis. J. Virol. 79: 2506–2516.

Stern-Ginossar, N. (2015). Decoding viral infection by ribosome profiling. J. Virol. 89: 6164–6166.

Stirrups, K., Shaw, K., Evans, S., Dalton, K., Casais, R., Cavanagh, D. and Britton, P. (2000). Expression of reporter genes from the defective RNA CD-61 of the coronavirus infectious bronchitis virus. J. Gen. Virol. 81:1687–1698.

Szklarczyk, D., Morris, J.H., Cook, H., Kuhn, M., Wyder, S., Simonovic, M., Santos, A., Doncheva, N.T., Roth, A., Bork, P., Jensen, L.J. and von Mering, C. (2017). The STRING database in 2017: quality-controlled protein-protein association networks, made broadly accessible. Nucleic Acids Res. 45(D1):D362–D368.

Tagliabracci, V.S., Wiley, S.E., Guo, X., Kinch, L.N., Durrant, E., Wen, J., Xiao, J., Cui, J., Nguyen, K.B., Engel, J.L., Coon, J.J., Grishin, N., Pinna, L.A., Pagliarini, D.J. and Dixon, J.E. (2015). A Single Kinase Generates the Majority of the Secreted Phosphoproteome. Cell 161:1619–1632.

Van Roekel, H., Clarke, M.K., Bullis, K.L., Olesiuk, O.M. and Sperling, F.G. (1951). Infectious bronchitis. Am. J. Vet. Res. 12:140–146.

Wolin, S.L. and Walter, P. (1988). Ribosome pausing and stacking during translation of a eukaryotic mRNA. EMBO J. 7: 3559–3569.

Zhang, J., Kaiser, M.G., Deist,. MS., Gallardo, R.A., Bunn, D.A., Kelly, T.R., Dekkers, J.C.M., Zhou, H.and Lamont, S.J. (2018). Transcriptome analysis in spleen reveals differential regulation of response to Newcastle disease virus in two chicken lines. Sci. Rep. 8:1278.

Zúñiga, S., Sola, I., Alonso, S. and Enjuanes, L. (2004). Sequence motifs involved in the regulation of discontinuous coronavirus subgenomic RNA synthesis. J. Virol. 78: 980–994.

